# Cellular uptake of folate-olaparib conjugates via folate receptor-mediated endocytosis: Potential for selective delivery of DNA damage response inhibitors into tumour cells

**DOI:** 10.64898/2026.09.23.752340

**Authors:** Vassilios Bavetsias, Lisa Pickard, Sara Diaz-Sanchez, Sharon Gowan, Francesca Wood, Ruth Ruddle, Christopher Wong, Humera H. Sharif, Florence I. Raynaud, Rosemary Burke, Anderson T. Wang, Julian Blagg, Vladimir Kirkin, Olivia W. Rossanese, Udai Banerji

## Abstract

The folate receptor α (FRα) is overexpressed in a range of human tumours including ovarian cancer cells. We propose that the overexpression of the FRα on the surface of ovarian tumour cells could be exploited for the selective delivery of a DNA damage response inhibitor (DDRi) in the form of an intact folate drug conjugate (FDC). This approach would improve the therapeutic index of the parent DDRi facilitating combination studies of the DDRi-based FDC with DNA damaging chemotherapy. FR-mediated cellular uptake of the proposed folate drug conjugates is requisite for FDC selective delivery into tumours. In this study, we synthesised a series of olaparib-based folate conjugates that maintained the biochemical PARP1 inhibition associated with olaparib and showed binding affinity for the folate receptor. Significantly, we identified compounds **10b** and **11** that selectively enter FRα overexpressing tumour cells via folate receptor-mediated endocytosis in their intact form and engage with their target as demonstrated by the potent inhibition of PARylation (KB cells, PARylation IC_50_ = 5.7 and 3.9 nM; respectively).

## Introduction

The folate receptors (FRs) exist in four isoforms, FRα, FRβ, FRγ, and FRδ with FRα, FRβ, and FRδ being attached to the cell membrane via glycosylphosphatidylinositol (GPI) linkage^1,2^. The expression of the FRα is limited in normal adult tissues but the FRα is overexpressed in a range of human malignancies including those of ovary, breast, and lung^3^. The overexpression of FRα in cancer cells has been exploited for the selective delivery of anticancer agents into tumours with approaches that include folate conjugates bearing a cleavable linker,^4,5^ ADCs,^6,7^ and small molecules that bind to the FR and selectively enter cells using the folate receptor endocytosis process. The latter approach is exemplified by CT900 (BGC945, ONX-0801), a thymidylate synthase inhibitor which was designed to selectively enter cells via α-folate receptor mediated endocytosis^8^. Possessing this dual pharmacology in one chemical entity, CT900 is selectively delivered to FRα overexpressing tumour cells and exerts its antitumour activity by inhibiting thymidylate synthase^8^. In a recent Phase I clinical trial of CT900 at the recommended dose of 12 mg/m^2^, 9/25 (36%) of patients with ovarian cancer whose ovarian cancers expressed medium or high expression of FRα achieved a radiological partial response^9^. It was therefore demonstrated, with CT900, that a FRα targeted small-molecule anticancer agent can be selectively delivered to FRα overexpressing tumours leading to a significant antitumour effect in the clinical setting. To expand on these findings, we are testing the hypothesis that small-molecule anticancer agents conjugated to folic acid (FA) using *non-cleavable* linkers could be selectively delivered to FRα-overexpressing cancer cells, in intact form, engaging with their target to elicit anti-tumour activity (Figure 1). This new class of conjugates is termed folate-drug conjugates (FDCs).

**Figure 1.**
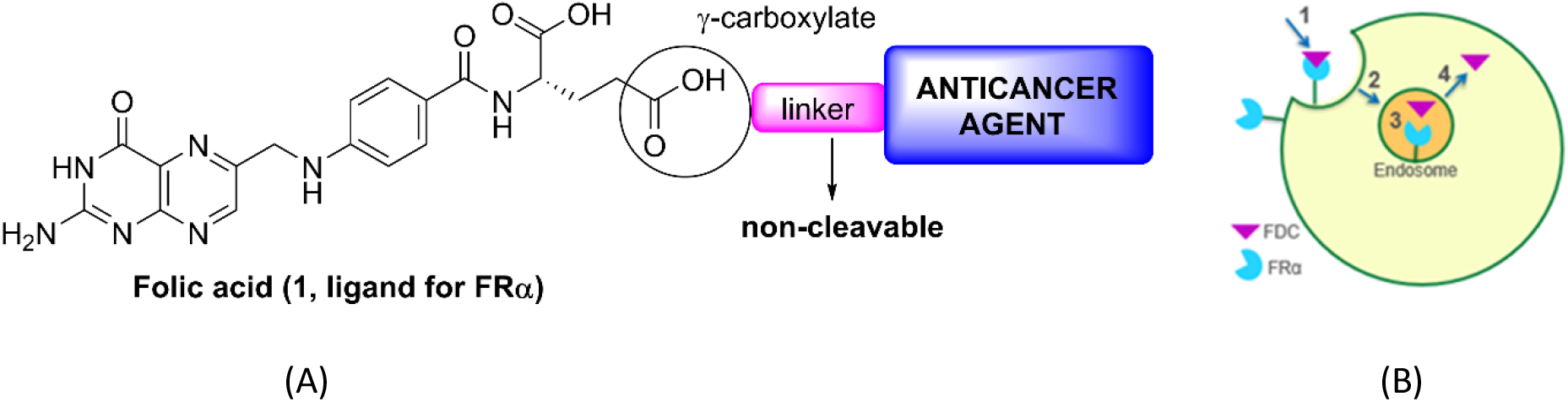
(A) Proposed FDCs for targeting FRα-overexpressing cancer cells. (B), Proposed mechanism for FDC cellular internalisation: 1. FDC binding to FRα, 2. endosome formation, 3. conjugate dissociation from the receptor by pH-driven conformational change, 4. conjugate escapes into cytosol and the FRα recycles onto the surface of the cell.

The main benefit of the FDC approach derives from the selective FDC delivery into the tumour cells that could lead to an improved therapeutic index of the parent anticancer agent. To this end, there have been an exciting range of DNA damage response (DDR) inhibitors evaluated in clinical trials with some being licensed for clinical use^10,11^. An exemplar of this class is olaparib (also known as AZD-2281, Lynparza) which is being used as a single agent in *BRCA*-mutant and platinum-sensitive ovarian cancer^10, 12^. Therapeutically, there is a strong rationale for combining a DNA damage response inhibitor (DDRi) with DNA damaging chemotherapy, and indeed it has been demonstrated that DDR inhibitors including olaparib sensitise cells to DNA damage chemotherapy^13,14^. However, this observed synergistic activity in preclinical models has not been translated into clinical benefit because of an enhanced normal tissue toxicity observed in clinical trials, in particular bone marrow toxicity, requiring reduction in the dose intensity of chemotherapy agents and olaparib itself ^15,16^. These findings led to the registration of olaparib as maintenance therapy after chemotherapy rather than in combination with chemotherapy in ovarian cancer^17,18^.

Taken all together, our therapeutic hypothesis is that selective delivery of a DDRi in the form of folate drug conjugate into FRα overexpressing tumour cells has the potential to significantly improve the therapeutic index of parent DDRi allowing combination studies of the DDRi-based FDC with DNA damaging chemotherapy. For this approach to succeed, it is prerequisite that the DDRi-based FDC selectively enters FRα overexpressing tumour cells via folate receptor mediated endocytosis. Herein, we present our work on addressing this key question by preparing and evaluating DDRi-derived FDCs based on the PARP1 inhibitor olaparib.

## Results and discussion

In the context of our hypothesis that selective delivery of a folate-DDRi conjugate into FRα overexpressing tumours would improve the therapeutic index of the parent DDRi, the main objective of this study was to show that olaparib-based FDCs bearing a *non-cleavable* linker could selectively enter cells via the folate receptor-mediated endocytosis process. The first step in the folate receptor-mediated cellular uptake involves binding to FR (Figure 1), thus binding affinity to the FR was investigated. This was followed by the determination of PARP1 inhibition in biochemical assays, we were aiming for olaparib FDCs to show PARP1 inhibitory activities similar to that of olaparib, a prerequisite in achieving potent target inhibition in cells. Additionally, FDC profiling included human plasma stability and Caco-2 permeability. In relation to the latter, we were aiming at low or lack of permeability in the non-FR Caco-2 cells since cellular uptake by passive diffusion would compromise selective delivery into tumour cells. We used two experimental approaches for demonstrating FR-mediated cellular uptake: i) we studied the inhibition of PARylation in FRα overexpressing cells (e.g. KB cells) and subsequently we investigated if any observed inhibition is reversed in the presence of folic acid, and ii) we studied the inhibition of PARylation in folate receptor negative cells (e.g. OV-90 cells) for those FDCs that potently inhibited PARylation in FRα overexpressing KB cells.

The design of olaparib-based FDCs was guided by the crystal structure of olaparib bound to PARP1 (pdb code: 5ds3)^19^ which shows the piperazine moiety pointing to solvent, suggesting the use of the piperazine nitrogen which is capped by the cyclopropyl acyl group in olaparib (Figure 2) as a suitable exit vector for linker attachment. It should be noted that successful piperazine derivatisation of olaparib in relation to PARP1 inhibition has been previously described for imaging in cells^20,21^.

**Figure 2.**
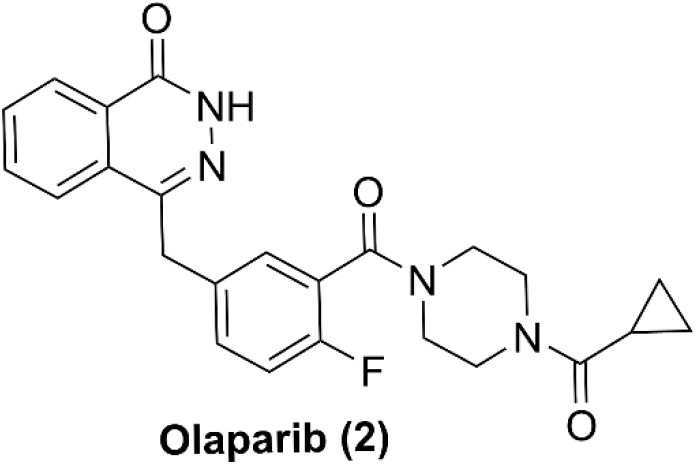
Chemical structure of olaparib.

Regarding conjugation to folic acid, we have chosen to preferentially attach the olaparib/linker moiety to the γ-glutamate carboxylate (Figures 1 and Tables 1, 2), an approach previously applied in the conjugation of various anticancer agents to folic acid via a cleavable linker with the resulting conjugates retaining affinity for the FRα and entering cells via FRα-mediated endocytosis^22,23^. The retention of affinity for FRα is rationalised by the crystal structure of folic acid bound to FRα that shows a solvent-exposed glutamate γ-carboxylate (pdb code: 4LRH)^24^.

**Table 1.**
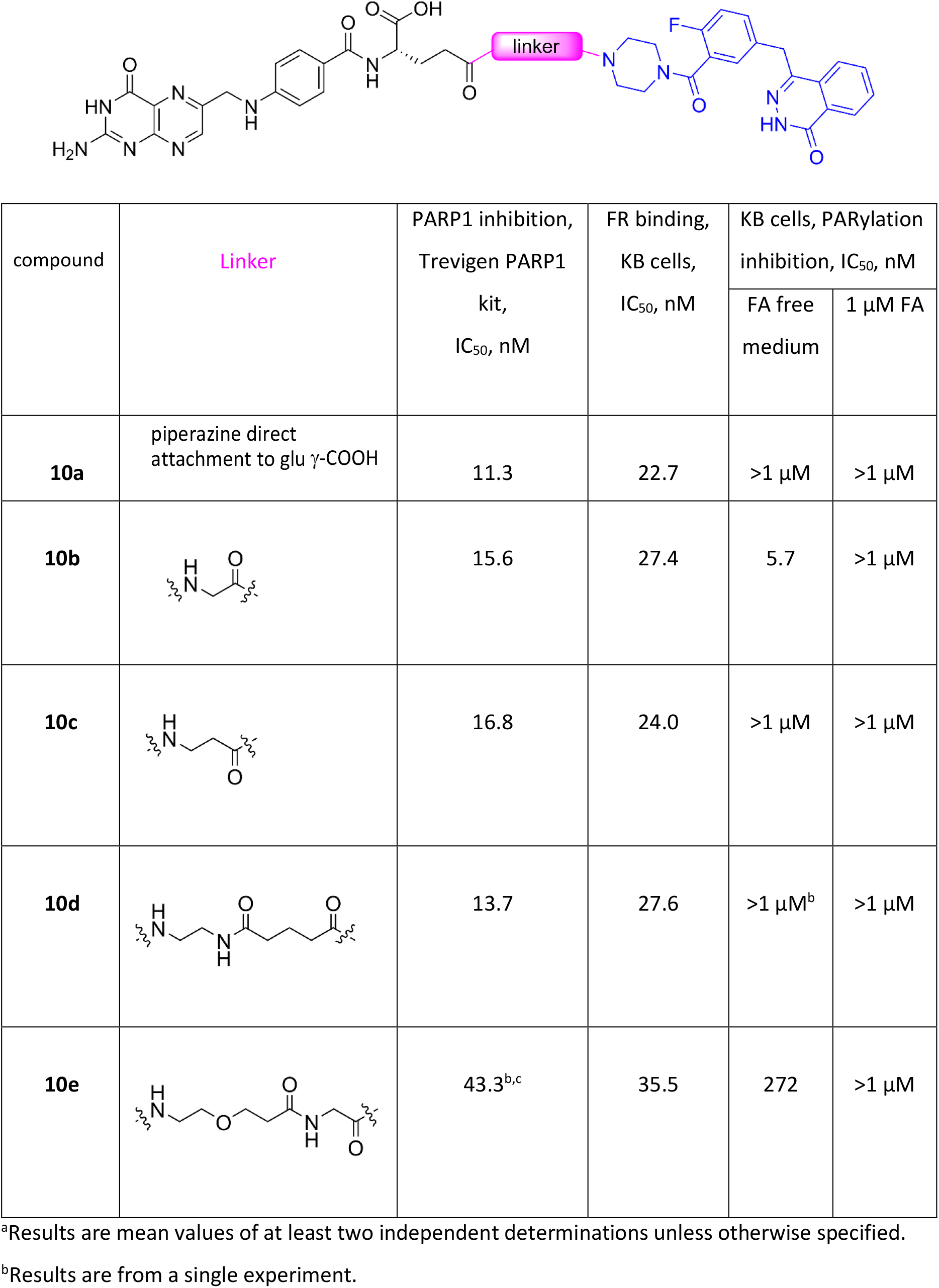
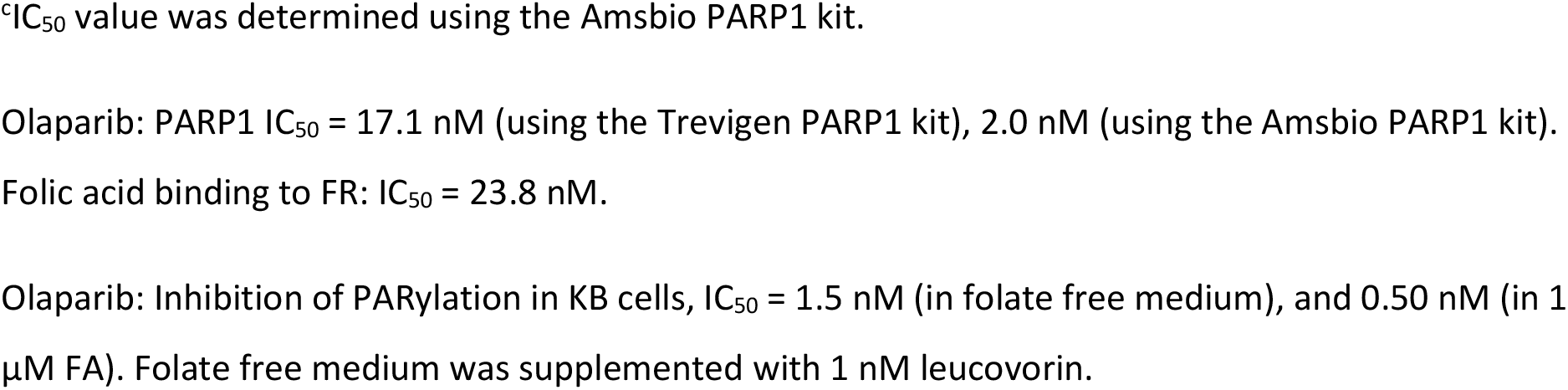
Olaparib-based FDCs, linker modifications: PARP1 inhibitiona, FR binding in KB cellsa, and inhibition of PARylation in KB cells, 48 ha.

| compound | Linker | PARP1 inhibition,<br>Trevigen PARP1<br>kit,<br>IC<sub>50</sub>, nM | FR binding,<br>KB cells,<br>IC<sub>50</sub>, nM | KB cells, PARylation<br>inhibition, IC<sub>50</sub>, nM |  |
| --- | --- | --- | --- | --- | --- |
| | | | | FA free<br>medium | 1 μM FA |
| **10a** | piperazine direct<br>attachment to glu γ-COOH | 11.3 | 22.7 | >1 μM | >1 μM |
| **10b** | | 15.6 | 27.4 | 5.7 | >1 μM |
| **10c** | | 16.8 | 24.0 | >1 μM | >1 μM |
| **10d** | | 13.7 | 27.6 | >1 μM<sup>b</sup> | >1 μM |
| **10e** | | 43.3<sup>b,c</sup> | 35.5 | 272 | >1 μM |

<sup>a</sup>Results are mean values of at least two independent determinations unless otherwise specified.
<sup>b</sup>Results are from a single experiment.
<sup>c</sup>IC<sub>50</sub> value was determined using the Amsbio PARP1 kit.
Olaparib: PARP1 IC<sub>50</sub> = 17.1 nM (using the Trevigen PARP1 kit), 2.0 nM (using the Amsbio PARP1 kit).
Folic acid binding to FR: IC<sub>50</sub> = 23.8 nM.
Olaparib: Inhibition of PARylation in KB cells, IC<sub>50</sub> = 1.5 nM (in folate free medium), and 0.50 nM (in 1 μM FA). Folate free medium was supplemented with 1 nM leucovorin.

**Table 2.**
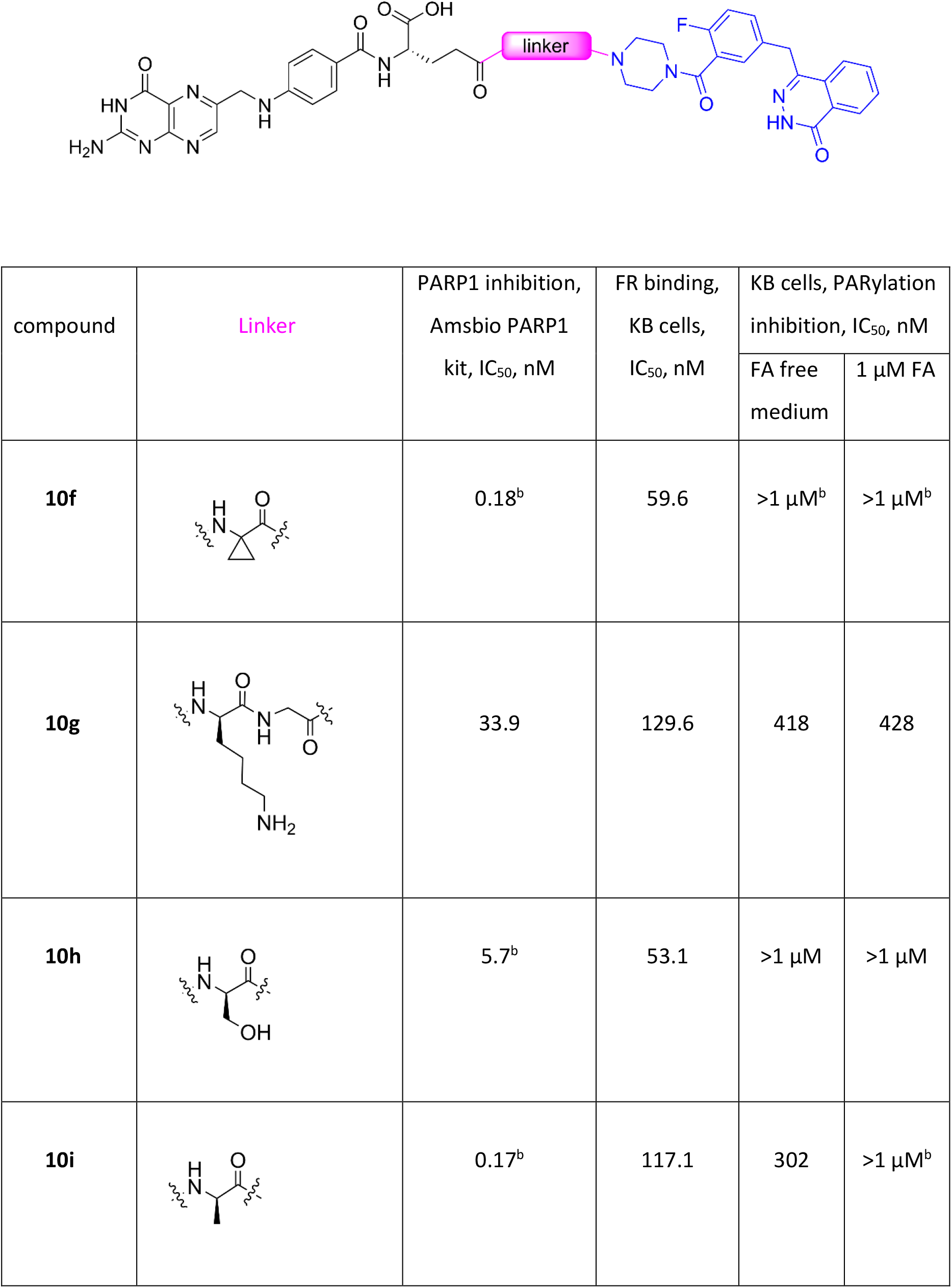
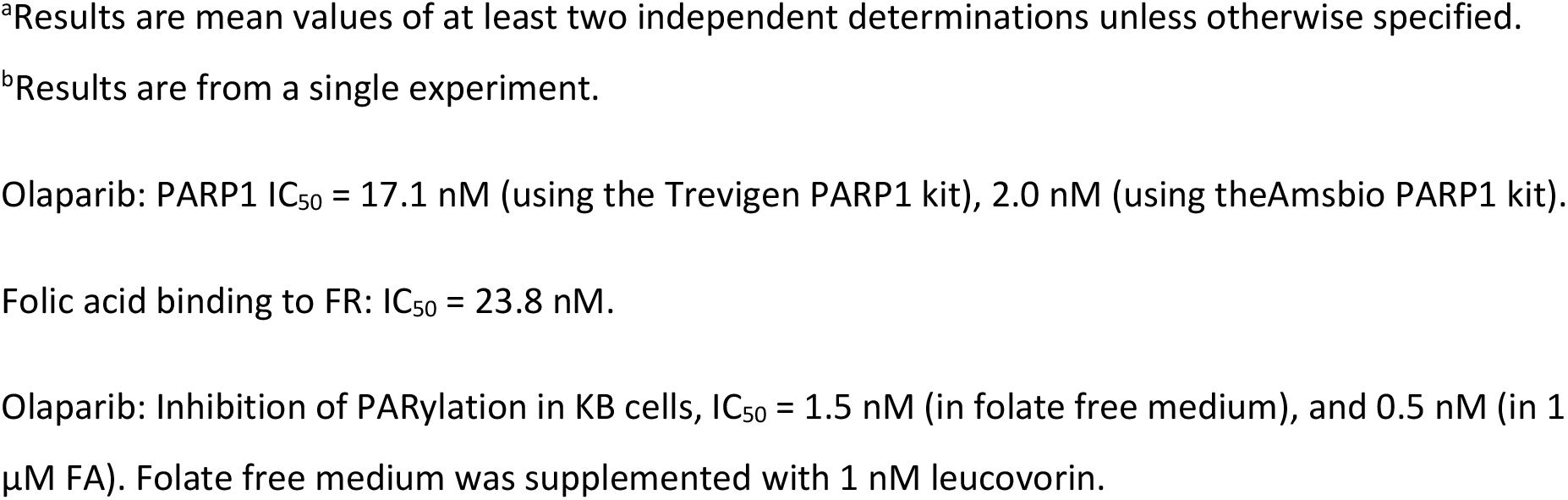
Olaparib-based FDCs, **10b** derivatives: PARP1 inhibition^a^, FR binding in KB cells^a^, and inhibition of PARylation in KB cells, 48 h^a^.

| compound | Linker | PARP1 inhibition,<br>Amsbio PARP1<br>kit, IC<sub>50</sub>, nM | FR binding,<br>KB cells,<br>IC<sub>50</sub>, nM | KB cells, PARylation<br>inhibition, IC<sub>50</sub>, nM |  |
| --- | --- | --- | --- | --- | --- |
| | | | | FA free<br>medium | 1 μM FA |
| **10f** | | 0.18<sup>b</sup> | 59.6 | >1 μM<sup>b</sup> | >1 μM<sup>b</sup> |
| **10g** |  | 33.9 | 129.6 | 418 | 428 |
| **10h** | | 5.7<sup>b</sup> | 53.1 | >1 μM | >1 μM |
| **10i** | | 0.17<sup>b</sup> | 117.1 | 302 | >1 μM<sup>b</sup> |

<sup>a</sup>Results are mean values of at least two independent determinations unless otherwise specified.
<sup>b</sup>Results are from a single experiment.
Olaparib: PARP1 $IC_{50}$ = 17.1 nM (using the Trevigen PARP1 kit), 2.0 nM (using the Amsbio PARP1 kit).
Folic acid binding to FR: $IC_{50}$ = 23.8 nM.
Olaparib: Inhibition of PARylation in KB cells, $IC_{50}$ = 1.5 nM (in folate free medium), and 0.5 nM (in 1 μM FA). Folate free medium was supplemented with 1 nM leucovorin.

**Scheme 1:**
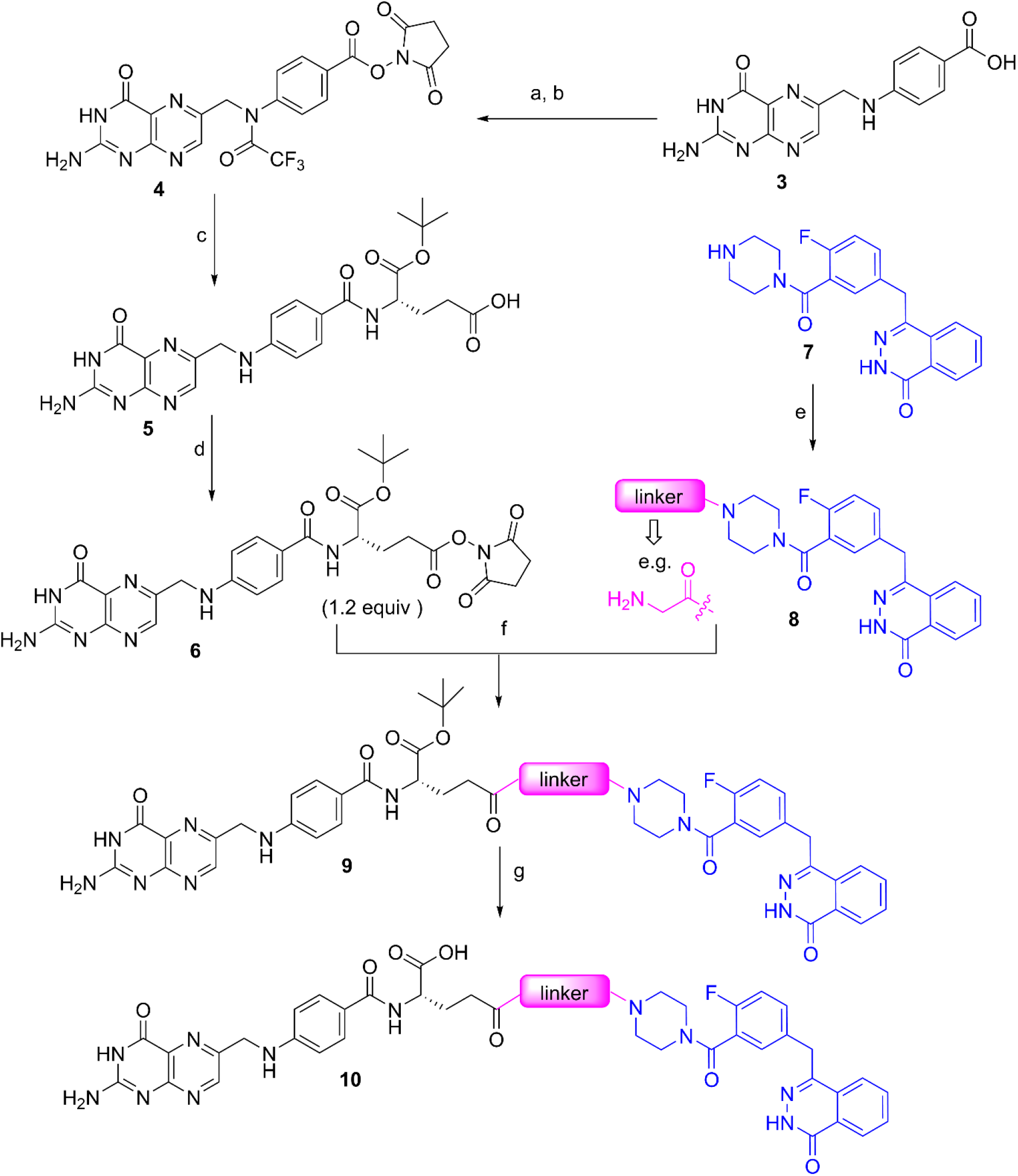
General route to olaparib FDCs. Reagents and conditions: (a) i) TFAA, ii) 3% TFA , aqueous NH_4_OH to neutralise; (b) NHS, DIC, DMAP, DMF; (c) i) L-glutamic acid α-*tert* butyl ester, Et_3_N, DMF, 18 h; ii) 7N NH_3_ in MeOH, room temp., 1 h; (d) NHS, DMAP, DIC, DMF, room temp., 40 h; (e) i) requisite linker, coupling reagent for amide bond formation, ii) removal of the amino protecting group; (f) Et_3_N, DMSO, room temp., 16 h in the dark; (g) TFA, room temp. 1 h, then concentration of the reaction mixture followed by purification.

Access to the olaparib FDCs **10a-e** and **10g-i** presented in Tables 1 and 2 was achieved by the route shown in Scheme 1 which involves the attachment of the linker to des-cyclopropylacyl olaparib (compound **7**) followed by the coupling of the olaparib/linker moiety **8** with *N*-hydroxysuccinimide active ester **6** and subsequent removal of the *tert*-butyl ester protecting group (Scheme 1). The *N*-hydroxysuccinimide active ester **6**, a key intermediate in this synthetic sequence, was prepared from pteroic acid (**3**) in four steps (Scheme 1). In a modification of the route outlined in Scheme 1, FDC **10f** was prepared by coupling of the olaparib/linker moiety **8f** to Fmoc-Glu-OBut followed by Fmoc deprotection, coupling to *N*^10^-(trifluoroacetamido)pteroic acid OSu active ester (**4**) and subsequent removal of the protecting groups. FDC **11** (Figure 4) was prepared by coupling the enantiomer of *N*-hydroxysuccinimide active ester **6** to the appropriate olaparib/linker moiety **8b**.

We first varied the length of the linker by both amino acid and ethylene glycol-based elongation to obtain conjugates **10a-e** (Table 1) which potently inhibited PARP1, suggesting tolerance to the length and chemical composition of the linker in the PARP1 inhibition biochemical assay. All folate-olaparib conjugates (**10a-e**) showed high binding affinity for the folate receptor, similar to that of folic acid (Table 1). Additionally, all analogues showed low permeability in the Caco-2 assay (A to B and B to A flux < 2×10^-6^ cm/s, no measurable efflux), a desirable outcome since cellular permeability by passive diffusion would compromise selective FDC delivery to tumour cells. Analogues **10a-d** were tested in human plasma demonstrating FDC stability over 1 h period. FDC **10b** was also confirmed to be stable (<25% loss) in human plasma over 24 h. Subsequently, to establish if any of these analogues use the folate receptor endocytosis process for cellular uptake, we investigated the inhibition of PARylation in FRα overexpressing KB cells both in folate free medium (supplemented with 1 nM leucovorin) and in the presence of 1 µM folic acid which competes with the FDC for binding to the FR, thus precluding FDC FR-mediated cellular uptake. FDCs **10a, 10c**, and **10d** showed no inhibition of PARylation suggestive of insufficient cellular uptake, whereas **10e** showed weak inhibition of PARylation (IC_50_ = 272 nM) which was reversed in the presence of 1 μM folic acid (Table 1). However, **10b**, displayed potent inhibition of PARylation in FRα overexpressing KB cells (IC_50_ = 5.7 nM) which was reversed in the presence of 1 µM folic acid (IC_50_ > 1 μM); Table 1, Figure 3). Additionally, **10b** showed weak inhibition of PARylation in FR-negative OV90 cells which was similar with that observed in KB cells in the presence of 1 μM folic acid (Figure 3). Taken together, this data suggests that **10b** selectively uses the folate receptor endocytosis process for cellular entry and potently inhibits its target in FRα overexpressing KB cells.

**Figure 3:**
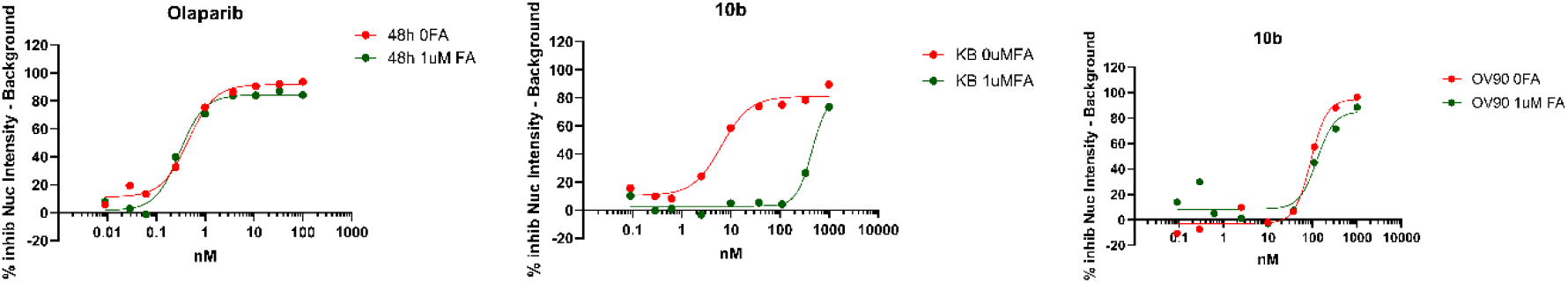
Left and middle panels: Inhibition of PARylation in FRα overexpressing KB cells in folate free medium (red) and in the presence of 1 µM folic acid (green) by olaparib and **10b**, respectively. Right panel: Inhibition of PARylation by **10b** in FR-negative OV90 cells in folate free medium (red) and in the presence of 1 µM folic acid (green).

**Figure 4:**
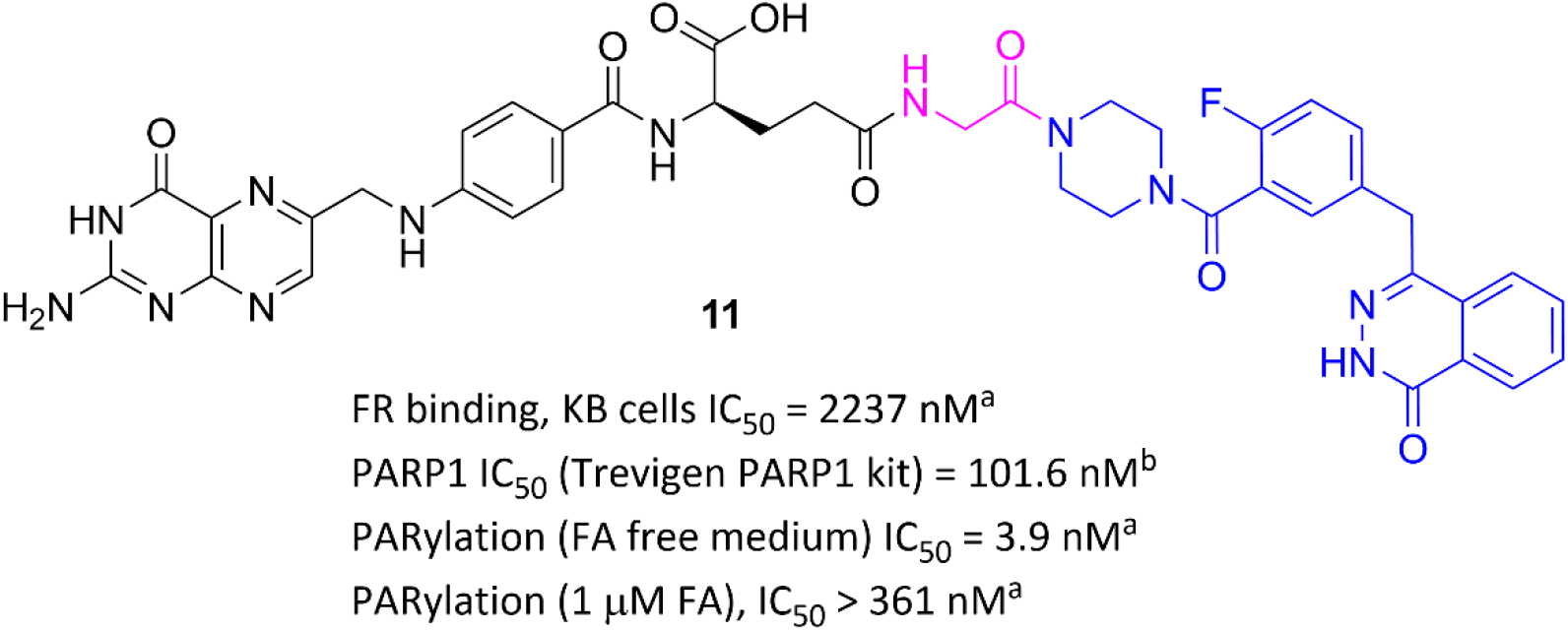
FDC **11** profiling. ^a^Results are mean values of at least two independent determinations.

Building on these findings, we continued linker variations based on the structure of **10b**, and we included in this exploration the introduction of hydrogen bond donor groups to further reduce the possibility of cellular uptake via passive diffusion.

The introduction of a cyclopropyl ring in the linker (FDC **10f**) was well tolerated in relation to PARP1 inhibition (biochemical assay) and binding to the FR (Table 2). FDC **10f** inhibited PARylation in FR overexpressing KB cells with IC_50_ value > 1 µM in both folate free medium and in the presence of folic acid (Table 2) which may suggest an insufficient cellular uptake. The introduction of a basic group into the linker in the form of a Lys side chain (FDC **10g**) was tolerated regarding PARP1 inhibition (biochemical assay) and binding to the FR (Table 2). Additionally, **10g** showed low permeability in the Caco-2 assay (A to B and B to A flux < 2×10^-6^ cm/s, no measurable efflux) and weak inhibition of PARylation in KB cells in both folate free medium and in the presence of 1 μM folic acid (Table 2). The D-serine counterpart of **10b** (FDC **10h**, Table 2) displayed potent inhibition of PARP1 in the biochemical assay and high binding affinity for the FR but showed no inhibition of PARylation in FRα overexpressing KB cells in both folate free medium and in the presence of folic acid (IC_50_ > 1 µM, Table 2), an indication of insufficient cellular uptake. On the other hand, the D-alanine counterpart of **10b** (FDC **10i**, Table 2) inhibited PARylation (IC_50_ = 302 nM, Table 2) in folate free medium with this inhibition weakened in the presence of 1 μM folic acid (IC_50_ > 1 µM, Table 2).

Next, we investigated the effect of chirality (Glu stereogenic centre) on FR binding and cellular uptake by FR-mediated endocytosis. To this end, the enantiomer of **10b** (FDC **11**, Figure 4) showed significant lower binding affinity for the folate receptor (IC_50_ = 2237 nM, Figure 4) compared to that observed with **10b** (IC_50_ = 27.4 nM, Table 1). However, **11** potently inhibited PARylation in FR overexpressing KB cells (IC_50_ = 3.9 nM) with this inhibition being reversed in the presence of 1 μM folic acid (IC_50_ > 361 nM) (Figures 4 and 5). Additionally, **11** showed a weak inhibition of PARylation in FR-negative OV90 cells (Figure 5). This data is consistent with **11** selectively using the FR-mediated endocytosis process for cellular uptake despite its weaker affinity for FR compared to **10b**.

**Figure 5:**
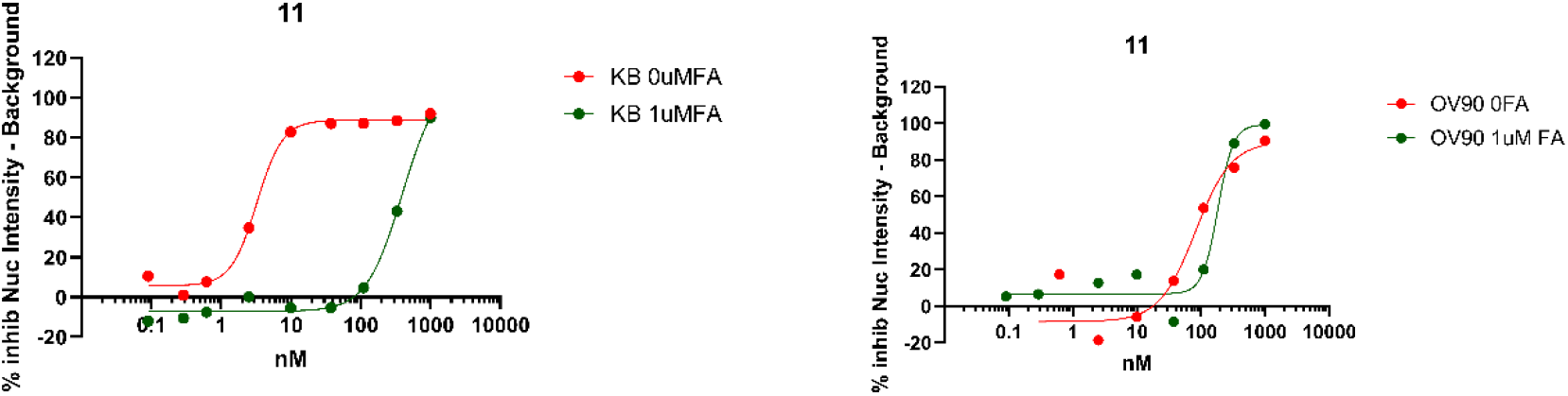
Left panel: Inhibition of PARylation by **11** in FRα overexpressing KB cells in folate free medium (red) and in the presence of 1 µM folic acid (green). Right panel: Inhibition of PARylation by **11** in FR-negative OV90 cells in folate free medium (red) and in the presence of 1 µM folic acid (green).

Having established **10b** and **11** as the most promising analogues for selective cellular uptake via folate receptor-mediated endocytosis and target engagement in cells (48 h incubation), we studied the effect of exposure time on inhibition of PARylation in KB cells at 2 h, 4 h, 6 h, 16 h, 24 h and 48 h. We observed that olaparib potently inhibits PARylation at 2 h (IC_50_ = 0.4 nM) with this inhibition being maintained up to 48 h (longest incubation period studied). Additionally, this inhibition was unaffected by co-incubation with 1 µM folic acid. These findings are consistent with a high cellular permeability by passive diffusion. On the other hand, we observed a delayed inhibition of PARylation by both **10b** and **11**, with no inhibition observed at 2 h. Significantly, **10b** and **11** exhibited distinctly different time dependent inhibitory profiles. FDC **11** potently inhibited PARylation at 6 h (IC_50_ = 12.1 nM), whereas a delayed inhibition was observed with **10b** which potently inhibited PARylation at 16 h with no significant inhibition observed at the earlier time points (2 h, 4 h, and 6 h) (Figure 6, Table 3). These findings suggest a more efficient trafficking of **11** through the cellular membrane. The profiling of **10b** and **11** reveals that their main difference relates to the binding affinity to the folate receptor, **11** is a much weaker binder of the folate receptor compared with **10b** (IC_50_ values of 2237 nM and 27.4 nM, respectively). The folate receptor mediated endocytosis is a multistep process as shown in Figure 1, and we hypothesise that the weaker binding of **11** to the folate receptor could facilitate dissociation of this FDC from the FR inside the endosome resulting in more efficient diffusion into cytosol and a faster intracellular FDC accumulation, consequently an earlier target engagement.

**Table 3.** Time dependent inhibition of PARylation by olaparib, **10b** and **11** in FRα overexpressing KB cells in folate free medium.^a^.

|  |  |  | **Olaparib** | **11** | **10b** |
| --- | --- | --- | --- | --- | --- |
| KB cells,<br>PARylation<br>inhibition<br>IC<sub>50</sub>, nM | 48 h | FA free media | 0.40 | 3.20 | 3.90 |
| | | 1 μM FA | 0.37 | 429.2 | 323.30 |
|  | 24 h | FA free media | 0.58 | 3.24 | 6.60 |
| | | 1 μM FA | 0.27 | 369.7 | 352.40 |
|  | 16 h | FA free media | 0.25 | 4.18 | 17.40 |
| | | 1 μM FA | 0.38 | 493.30 | 597.30 |
|  | 6 h | FA free media | 0.34 | 12.1 | 540.65 |
| | | 1 μM FA | 0.34 | >1000 | 936.70 |
|  | 4 h | FA free media | 0.70<sup>b</sup> | >1000<sup>b</sup> | >1000<sup>b</sup> |
| | | 1 μM FA | 0.83<sup>b</sup> | >1000<sup>b</sup> | >1000<sup>b</sup> |
|  | 2 h | FA free media | 0.41 | >1000 | >1000 |
| | | 1 μM FA | 0.35 | >1000 | >1000 |

<sup>a</sup>Results are mean values of at least two independent determinations unless otherwise specified.
<sup>b</sup>Results are from a single experiment.

**Figure 6:**
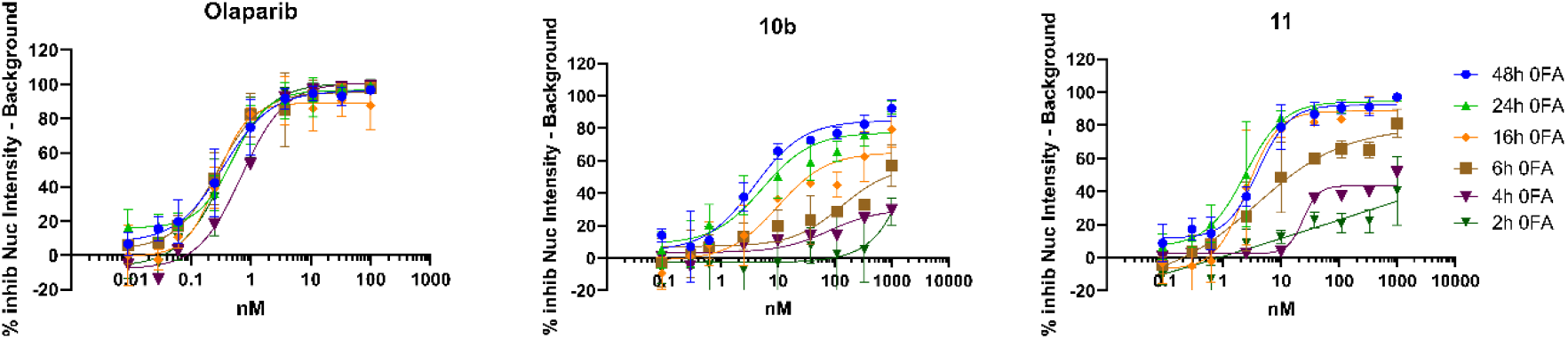
Time dependent inhibition of PARylation by olaparib, **10b** and **11** in FRα overexpressing KB cells. All time course experiments have been conducted in folate free medium and repeated at least 3 times (exception at 4 h: n=1).

## Conclusions

We synthesised a range of olaparib-folate conjugates by stable linker modifications and explored SAR trends towards inhibition of PARP1 in a biochemical assay, folate receptor binding and, importantly, cellular uptake via FR-mediated endocytosis. In general, the olaparib-based FDCs maintained biochemical PARP1 inhibition associated with olaparib and showed high binding affinity for the folate receptor. A significant drop in folate receptor binding affinity was observed when the glutamate in folic acid was replaced with its enantiomeric form (**10b** vs **11**). Cellular uptake via folate receptor-mediated endocytosis was studied by determining inhibition of PARylation in FRα overexpressing KB cells in folate free medium and in co-incubation with 1 µM folic acid. The majority of FDCs showed no inhibition of PARylation indicating insufficient cellular uptake; however, we identified a small number of FDCs using the folate receptor for cellular uptake, notably **10b** and **11**. These results support the hypothesis of targeting the folate receptor for the selective delivery of an intact DDRi-based FDC into FRα overexpressing tumour cells. The selective delivery of DDRi-based FDCs in tumours should improve the therapeutic index of the parent DDRi facilitating combination therapy with DNA damaging anticancer agents in FRα overexpressing tumours such as ovarian cancer.

## Materials and Methods

### PARP1 inhibition (biochemical assay)

Olaparib and olaparib-FDCs activity has been screened using the HT Homogeneous PARP Inhibition Assay Kit from Trevigen® (Cat#4690-096-K). The conjugates are incubated with 35nM PARP enzyme for a 15 mins and then Cycling enzymes (detection reaction) are added in a final volume of 10µL in a 384 well plate. The fluorescence is detected during detection reaction for 60 min (1 read per minute) using PheraStar (Ex./Em. 544/590nm).

### PARP1 inhibition (PARP1 chemiluminescent assay)

Due to the above kits being discontinued, Olaparib and olaparib-FDCs activity have also been screened using the Chemiluminescent assay from Amsbio (Cat# AMS.80551). In this protocol, 96 well plates were coated with Histone overnight at 4°C. The conjugates were then incubated with a master mix of 10X Parp buffer, 50ng PARP enzyme, 10x assay mixture and Activated DNA (5x) for 60 mins. Detection proceeded with the addition of streptavidin-HRP for 30 minutes in a final volume of 50µL and the signal generated by addition of HRP chemiluminescent substrates (A and B) (50ul each). Chemiluminescence was detected immediately using Victor plate reader.

### Folate receptor binding

The affinity of the olaparib-FDCs for folate receptor has been assessed by competition binding in 384 well plate using fluorescent folic acid labelled (FR680) from Perkin Elmer (#NEV10040). KB cells are pre-treated with unlabelled folic acid or olaparib-based FDCs at different concentrations (0-6µM) for 10 mins then 1µM FR680 is added to the cells and incubated for 1h at 37°C. The amount of the conjugate or folic acid bound to the folate receptor is determined by IF using INCELL2200.

### PARP1 inhibition in cells (PARylation assay)

FDC target engagement assay in a 96 well plate-based format. KB cells in folic acid free media were pre-treated with +/- Folic acid (1µM, 10mins) followed by 48h incubation with FDC. Olaparib is added as a control. DNA damage is induced by adding 10mM hydrogen peroxide for the last 10 mins. Inhibition of PARP results in a decrease in PAR expression which was quantified using an anti-PAR monoclonal antibody (4335-MC-100) by immunofluorescent imaging.

### Compound synthesis

Experimental details for the synthesis of key intermediates and final compounds are provided in electronic supplementary information.

## Author contribution

V. B. conceptualisation, funding acquisition, supervision, writing – original draft, writing – review & editing; L. P. investigation, data curation; S. D-S. investigation, data curation; S. G. investigation, data curation; F. W. investigation, data curation; R. R. investigation, data curation; C. W. investigation, data curation; H. H. S. investigation, data curation; F. I. R. supervision; R. B. supervision; A. T. W. supervision, methodology, writing – review & editing; J. B. conceptualisation, funding acquisition, writing – review & editing; V. K. supervision, project administration; O. W. R. supervision; U. B. conceptualisation, funding acquisition, supervision, project administration, writing – review & editing.

## Acknowledgments

This research was supported by Cancer Research UK [grant numbers C2739/A22897 and C309/A31322] and ICR MRC Confidence in Concept (CiC) grants (MC_PC_17163 and MC_PC_18051). The authors would like to thank Meirion Richards, Maggie Liu, and Amin Mirza of the Structural Chemistry Team within the Centre for Cancer Drug Discovery at the Institute of Cancer Research for their expertise and assistance. Authors acknowledge infrastructural funding to The Institute of Cancer Research and the Royal Marsden Hospital Foundation Trust for the Experimental Cancer Medicine Centre and the Biomedical Research Centre grants. UB, LP and RR acknowledge infrastructural funding from the Cancer Research Convergence Science Centre.

