## Supplementary Information for "Cellular uptake of folate-olaparib conjugates via folate receptor-mediated endocytosis: Potential for selective delivery of DNA damage response inhibitors into tumour cells"

**Table of contents**

Experimental details for the synthesis of key intermediates and final compounds .........pages 2-28

**Compound synthesis – General Experimental**

NMR: Routine $^1$H NMR spectra were recorded on a Bruker 500 MHz or 400 MHz spectrometer. Where stated, chemical shifts are expressed in parts per million values (ppm) and are designated as s (singlet); br s (broad singlet); d (doublet); dd (double doublet); ddd (double double doublet); dt (double triplet); t (triplet); br t (broad triplet); q (quartet); quint (quintet) or m (multiplet).

Methods for LC-MS analysis:

**Method A:** Waters Acquity H class UPLC with diode array (210-350 nm) and QDa mass detector. Column: Acquity UPLC CSH C18 1.7 µm 2.1 x 50 mm (Flow 0.6 mL/min). Run Time: 3.00 min. Conditions: Water [eluent A], MeCN [eluent B], 0.05% Formic Acid in water [eluent C]. Hold 90% A/5% B/5%C 1.80 min, gradient: 5-95% B/5% C in 0.4 min, hold at 95% B/5%C to 2.30 min.

**Method B:** Waters Acquity H class UPLC with diode array (210-350 nm) and QDa mass detector. Column: Acquity UPLC BEH C18 1.7 µm 2.1 x 50 mm (Flow 0.6 mL/min). Run Time: 3.00 min. Conditions: Water [eluent A], MeCN [eluent B], 0.05% Ammonia in water [eluent C]. Hold 90% A/5% B/5%C to 1.80 min, gradient: 5-95% B/5% C in 0.4 min, hold at 95% B/5%C to 2.30 min.

**Method C:** Waters Alliance HT Separations module 2790 with diode array (210-350 nm) and waters micromass ZQ. Column: YMC-Triart C18 1.7 µm 2.0 x 50 mm (Flow 0.8 mL/min). Run Time: 5.00 min. Conditions: Water [eluent A], MeCN [eluent B], 1% Formic in water:MeCN 1:1 [eluent C]. Gradient 0.0 - 4.0 mins 0-95% B, 5% C; 4.0-4.4 mins 95% B, 5% C; 4.4-4.5 mins 95% A, 5% B. Hold 95% A, 5% B to 5 min.

**Method D:** Waters Alliance HT Separations module 2790 with diode array (210-350 nm) and waters micromass ZQ. Column: YMC-Triart C18 1.7 µm 2.0 x 50 mm (Flow 0.8 mL/min). Run Time: 5.00 min. Conditions: Water [eluent A], MeCN [eluent B], 1% Ammonia in water:MeCN 1:1 [eluent D]. Gradient: 0.0 - 4.0 mins 0-95% B, 5% D; 4.0-4.4 mins 95% B, 5% D; 4.4-4.5 mins 95% A, 5% B. Hold 95% A, 5% B to 5 min.

**Method E:** Mass spectra were run on LC-MS systems using electrospray ionization. These were run using a Waters Acquity H-Class UPLC with PDA and QDa mass detection. Column: Acquity UPLC BEH C18 2.1x100mm 1.7µm. Column Temp 50 ˚C. Eluents: A: H20, 0.1% ammonia B: acetonitrile. Flow Rate: 0.6 mL/min. Gradient: 0.5-6.5mins 2-98%B 6.5-7.5mins 98% B.

**Method F:** Mass spectra were run on LC-MS systems using electrospray ionization. These were run using a Waters Acquity H-Class UPLC with PDA and QDa mass detection. Column: Acquity UPLC BEH C18 2.1 x 50 mm 1.7 µm. Column Temp: 50 ˚C. Eluents: A: $H_2O$, 0.1% ammonia B: MeCN. Flow Rate: 0.6 mL/min. Gradient: 0.2-2.5 min 2-98% B, 2.5-3.3 min 98% B, 3.3-3.5 98% A.

**Method G:** Mass spectra were run on LC-MS systems using electrospray ionization. These were run using a Waters Acquity H-Class UPLC with PDA and QDa mass detection. Column: Acquity UPLC BEH C18 2.1 x 50 mm 1.7 µm. Column Temp: 50 ˚C. Eluents: A: $H_2O$, 0.1% formic acid B: MeCN. Flow Rate: 0.6 mL/min. Gradient: 0.2-2.5 min 2-98% B, 2.5-3.3 min 98% B, 3.3-3.5 98% A.

**Method H:** Mass spectra were run on LC-MS systems using electrospray ionization. These were run using a Waters Acquity H-Class UPLC with PDA and QDa mass detection. Column: Acquity UPLC BEH C18 2.1 x 50 mm 1.7 µm. Column Temp: 50 ˚C. Eluents: A: $H_2O$, B: MeCN. Gradient: 0.00-1.70 min 0 - 95% B, 1.70 – 2.10 min 95% B, 2.10 – 2.50 min 95% A.

**Method I:** Mass spectra were run on LC-MS systems using electrospray ionization. These were run using a Waters Acquity H-Class UPLC with PDA and QDa mass detection. Column: Acquity UPLC BEH C18 2.1 x 50 mm 1.7 µm. Column Temp: 50 ˚C. Eluents: A: $H_2O$, B: MeCN. Gradient: 0.00-0.5 min 95% A, 0.50 – 6.50 min 0-95% B, 6.50 – 7.50 min 95% B, 7.50 – 8.00 95% A.

**Synthesis of key intermediate 6**

**4-(N-((2-amino-4-oxo-3,4-dihydropteridin-6-yl)methyl)-2,2,2-trifluoroacetamido)benzoic acid**

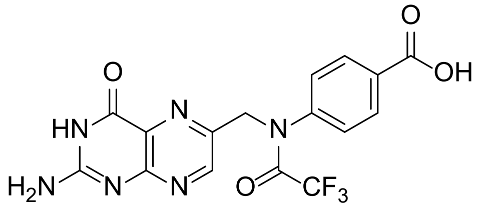


To a vigorously stirred solution of trifluoroacetic anhydride (76.0 mL, 546.0 mmol) in a 250 mL round bottom flask was added, portion wise, pteroic acid (3.28 g, 10.50 mmol) and the reaction mixture was stirred at room temperature for 16 h. The reaction mixture was then concentrated to a brown viscous oil/solid residue was suspended in THF (98 mL) which followed by TFA (0.33 mL) and ice (6.6 g). The reaction mixture was stirred at room temperature for 3 h and combined with material obtained from previous experiment performed at 1 g (3.2 mmol scale). Diethyl ether (650 mL) was then added, and the mixture was stirred at room temperature for 20 min. The precipitate was collected by filtration and washed with diethyl ether (2 x 30 mL) to afford the title compound as a brown solid (4.85 g, 11.88 mmol, 87%).

LCMS (Method C): RT = 1.59 min, ES+ 409.0 $[M+H]^+$.

**2,5-dioxopyrrolidin-1-yl4-(N-((2-amino-4-oxo-3,4-dihydropteridin-6-yl)methyl)-2,2,2-trifluoroacetamido)benzoate (4)**

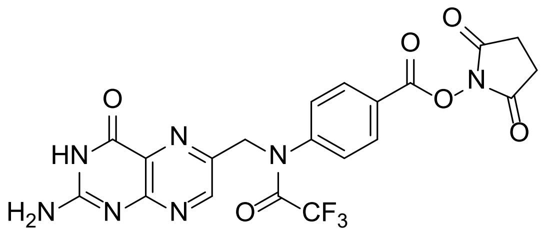


To a stirred solution of 4-(N-((2-amino-4-oxo-3,4-dihydropteridin-6-yl)methyl)-2,2,2-trifluoroacetamido)benzoic acid (4.85 g, 11.9 mmol) , NHS (2.74 g, 23.8 mmol) and DMAP (73 mg, 0.6 mmol) in anhydrous DMF (121 mL) was added DIC (2.8 mL, 17.9 mmol), and stirring was continued at room temperature for 64 h in the dark. Concentration of the solution in vacuo afforded a dark brown solid which stirred in DCM (120 mL) for 30 min. The precipitate was collected by filtration and washed with dichloromethane (2 x 100 mL) and diethyl ether (2 x 50 mL) to give a brown solid (5.21 g) which was further washed with dichloromethane (2 x 50 mL) and diethyl ether (2 x 50 mL) to afford the title compound as a brown solid (4.60 g, 76%).

LCMS (Method C): RT = 1.86 min, ES+ 506.1 $[M+H]^+$.

**(*S*)-4-(4-(((2-amino-4-oxo-3,4-dihydropteridin-6-yl)methyl)amino)benzamido)-5-(*tert*-butoxy)-5-oxopentanoic acid (5)**

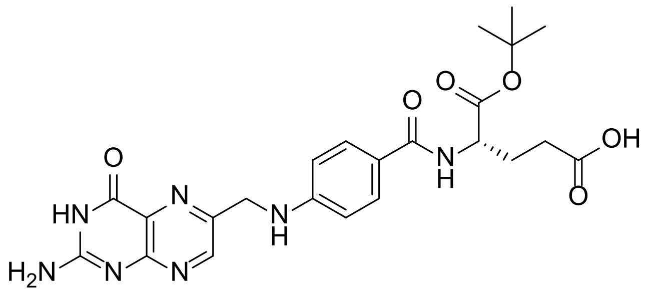

To a stirred mixture of 2,5-dioxopyrrolidin-1-yl 4-(*N*-((2-amino-4-oxo-3,4-dihydropteridin-6-yl)methyl)-2,2,2-trifluoroacetamido)benzoate (3.5 g, 6.93 mmol) in anhydrous DMF (87.5 mL) was added L-Glu-O$^t$Bu (2.83 g, 13.9 mmol) and triethylamine (2.0 mL, 13.9 mmol), and stirring was continued for 2 h in the dark. Ammonia in methanol (7N; 35 mL) was added and stirring was continued at room temperature for 1h. The mixture was then concentrated in vacuo and the residue was suspended in water (200 mL) and stirred at room temperature for 30 min. The precipitate was collected by filtration, washed with water (3 x 100 mL) and dried under vacuum overnight to afford the title compound as a brown solid (2.54 g, 74%).

$^1$H NMR (500 MHz, DMSO-$d_6$) δ 8.64 (s, 1H), 8.21 (brs, 1H), 7.65 (d, *J* = 8.7 Hz, 2H), 6.93 (t, *J* = 5.9 Hz, 1H), 6.88 (br s, 2H), 6.64 (d, *J* = 8.7 Hz, 2H), 4.48 (d, *J* = 5.9 Hz, 2H), 4.29-4.23 (m, 1H), 2.31 (t, *J* = 7.6 Hz, 2H), 2.05-1.95 (m, 1H), 1.93-1.84 (m, 1H), 1.39 (s, 9H).

LCMS (Method C): RT = 1.64 min, ES+ 498.2 $[M+H]^+$.

**1-(*tert*-butyl)5-(2,5-dioxopyrrolidin-1-yl)(4-(((2-amino-4-oxo-3,4-dihydropteridin-6-yl)methyl)amino)benzoyl)-*L*-glutamate (6)**

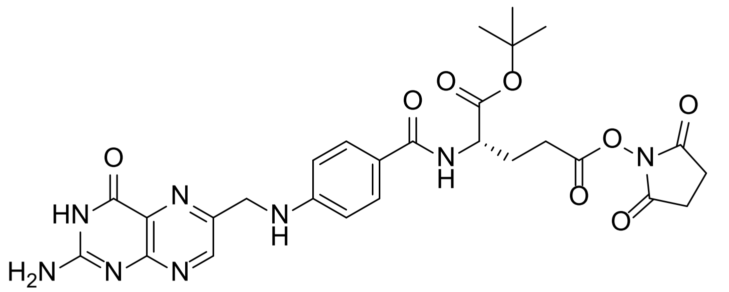

In a cooled mixture of (S)-4-(4-(((2-amino-4-oxo-3,4-dihydropteridin-6-yl)methyl)amino)benzamido)-5-(*tert*-butoxy)-5-oxopentanoic acid (**5**) (2.5 g, 5.02 mmol), NHS (1.74 g, 15.1 mmol), DMAP (31 mg,

0.25 mmol) and anhydrous DMF (78 mL) at 0 °C was added dropwise DIC (1.6 mL, 10.04 mmol). The stirred reaction mixture was allowed to warm to room temperature and stirring was continued at this temperature for 40 h in the dark. The mixture was then concentrated to a black/brown gum which was treated with dichloromethane / diethyl ether (v/v 3:1; 120 mL) and stirred at room temperature for 30 min. The precipitate was collected by filtration and washed with diethyl ether (2 x 30 mL) and then suspended in dichloromethane (100 mL). The precipitate was collected by filtration and washed with diethyl ether (2 x 50 mL) and dried to afford the title compound as a brown solid in quantitative yield.

$^1$H NMR (500 MHz, DMSO-$d_6$) δ 11.47 (br s, 1H), 8.65 (s, 1H), 8.21 (d, *J* = 7.8 Hz, 1H), 7.66 (d, *J* = 8.7 Hz, 2H), 6.96 (t, *J* = 6.0 Hz, 1H), 6.89 (brs, 2H), 6.65 (d, *J* = 8.7 Hz, 2H), 4.49 (d, *J* = 6.0 Hz, 2H), 4.35-4.29 (m, 1H), 2.81 (s, 4H), 2.89 – 2.72 (m, 2H), 2.16 – 1.99 (m, 2H), 1.39 (s, 9H).

LCMS (Method C): RT = 1.90 min, ES+ 595.3 $[M+H]^+$.

**Synthesis of 10a**

**(S)-2-(4-(((2-amino-4-oxo-3,4-dihydropteridin-6-yl)methyl)amino)benzamido)-5-(4-(2-fluoro-5-((4-oxo-3,4-dihydrophthalazin-1-yl)methyl)benzoyl)piperazin-1-yl)-5-oxopentanoic acid (10a)**

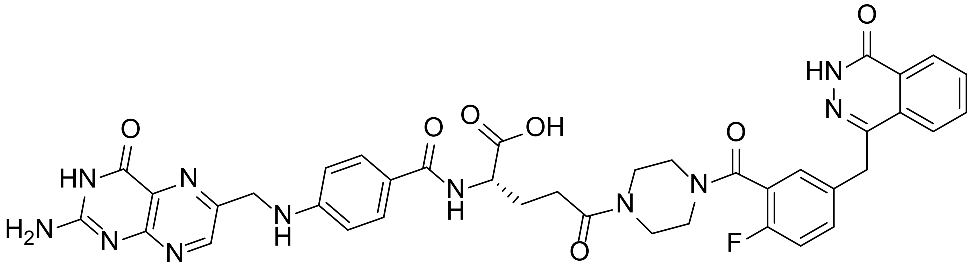

To a stirred mixture of amine **7** (22 mg, 0.06 mmol) and folate active ester **6** (40 mg, 0.07 mmol) in DMSO (1 mL) was added triethylamine (43 mg, 58 µL, 0.42 mmol) and stirring was continued at room temperature in the dark for 64 h. Purification by reversed-phase chromatography (Combiflash, C18 20 g, 2 → 45 % $CH_3CN$ in 0.1 % aq. $NH_3$) gave the intermediate *tert*-butyl ester product as a brown solid after concentration. The intermediate was immediately stirred in TFA (3 mL) for 1 h. The mixture was concentrated and 7 N $NH_3$ in MeOH (0.5 mL) and 0.1 % aqueous ammonia (1.5 mL) were added. Purification by reversed-phase chromatography (Combiflash, C18 20 g, 2 → 40 % $CH_3CN$ in 0.1 % aq. $NH_3$) gave the title compound as a pale brown solid (14 mg, 30 %) after freeze-drying.

$^{1}H$ NMR (500 MHz, DMSO-$d_6$) δ 12.58 (2 x s, 1H), 8.63 (s, 1H), 8.25 (d, $J$ = 7.7 Hz, 1H), 7.99 – 7.85 (m, 3H), 7.85 – 7.77 (m, 1H), 7.60 (t, 2H), 7.44 – 7.40 (m, 1H), 7.35 (dd, $J$ = 14.0, 6.5 Hz, 1H), 7.22 (t, $J$ = 8.9 Hz, 1H), 7.03 (br s, 2H), 6.93 – 6.88 (m, 1H), 6.64 (t, 2H), 4.50 – 4.44 (m, 2H), 4.32 (s, 2H), 4.19 – 4.15 (m, 1H), 3.61 – 3.58 (m, 2H), 3.50 – 3.45 (m, 2H), 3.34 (s, overlap with $H_2O$), 3.15 – 3.11 (m, 2H), 2.33 (m, 2H), 2.06 – 2.01 (m, 1H), 1.98 – 1.86 (m, 1H).

LCMS (Method B): RT = 0.96 min, >98 % @ 254 nm, ES+ 790.2 $[M+H]^+$.

HRMS: found 790.2856; calculated for $C_{39}H_{37}FN_{11}O_7$ $(M+H)^+$ 790.2861.

**Synthesis of 10b**

**4-({4-Fluoro-3-[(4-glycyl-1-piperazinyl)carbonyl]phenyl}methyl)-2*H*-phthalazin-1-one Trifluoroacetate (8b)**

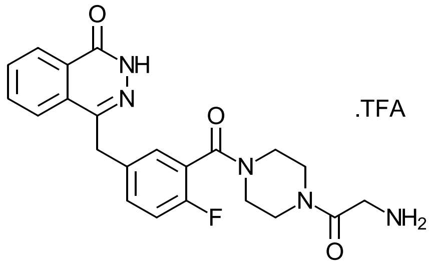


To a stirred solution of 4-({4-fluoro-3-[(1-piperazinyl)carbonyl]phenyl}methyl)-2*H*-phthalazin-1-one (**7**) (200 mg, 0.55 mmol), *N*-Boc-glycine (107 mg, 0.61 mmol) and DIPEA (142 mg, 191 µL, 1.1 mmol) in DMF (5 mL) was added HATU (105 mg, 0.55 mmol) and stirring was continued at room temperature for 18 h. Analysis by LCMS showed ~70 % conversion. *N*-Boc-glycine (49 mg, 0.28 mmol) and HATU (54 mg, 0.28 mmol) were added and stirring continued for 1 h giving > 90 % conversion by LCMS. The reaction mixture was diluted with ethyl acetate (40 mL) and washed with water (40 mL). The aqueous layer was back extracted with ethyl acetate (40 mL). Combined organic layers were washed with water (60 mL), 2 M hydrochloric acid (60 mL), water (60 mL), sat. aqueous $NaHCO_3$ (60 mL), brine (60 mL), dried ($MgSO_4$) and concentrated to give the Boc-protected product as a beige solid (220 mg, 76 %).

LCMS (Method C): RT = 2.35 min, ES+ 524.2 $[M+H]^+$.

A solution of the Boc-protected product (117 mg, 0.22 mmol) was stirred in TFA (1 mL) at room temperature for 1 h. The mixture was concentrated and residual TFA removed by azeotrope with dichloromethane (3 × 3 mL). The resulting gum was triturated with diethyl ether (2 × 3 mL, decanted off) and dried to a colourless gum (125 mg, quant.) that was used directly without further purification.

$^{1}H$ NMR (500 MHz, DMSO-$d_6$) δ 12.60 (s, 1H), 8.27 (d, *J* = 7.9 Hz, 1H), 8.07 – 8.01 (m, 3H), 7.99 – 7.93 (m, 1H), 7.93 – 7.86 (m, 1H), 7.83 (t, *J* = 7.6 Hz, 1H), 7.50 – 7.44 (m, 1H), 7.36 (dd, *J* = 16.4, 6.3 Hz, 1H), 7.25 (t, *J* = 9.0 Hz, 1H), 4.34 (s, 2H), 3.92 (q, *J* = 5.7 Hz, 1H), 3.85 (q, *J* = 5.6 Hz, 1H), 3.68 – 3.57 (m, 3H), 3.44 (dt, *J* = 16.7, 5.4 Hz, 2H), 3.33 – 3.28 (m, 1H), 3.26 – 3.21 (m, 1H), 3.21 – 3.15 (m, 1H).

LCMS (Method D): RT = 1.78 min, ES+ 424.2 $[M+H]^+$.

**$N^2$-(4-(((2-amino-4-oxo-3,4-dihydropteridin-6-yl)methyl)amino)benzoyl)-$N^5$-(2-(4-(2-fluoro-5-((4-oxo-3,4-dihydrophthalazin-1-yl)methyl)benzoyl)piperazin-1-yl)-2-oxoethyl)-*L*-glutamine (10b)**

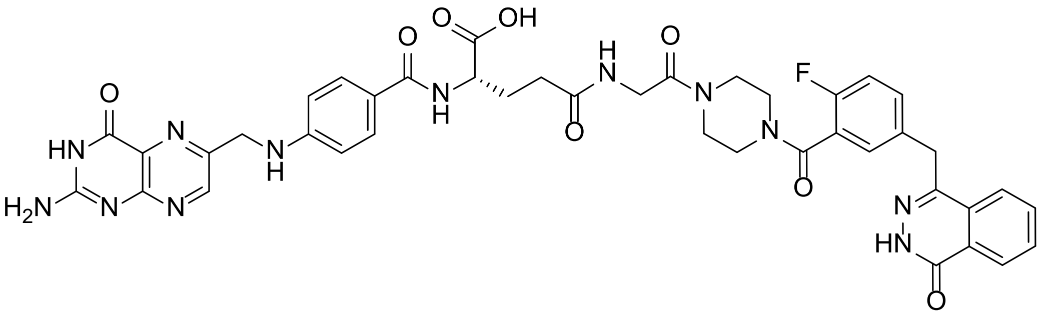


To a stirred mixture of 4-({4-fluoro-3-[(4-glycyl-1-piperazinyl)carbonyl]phenyl}methyl)-2*H*-phthalazin-1-one trifluoroacetate (**8b**) (26 mg, 0.05 mmol) and the folate active ester **6** (31 mg, 0.06 mmol) in DMSO (1 mL) was added triethylamine (34 mg, 47 µL, 0.34 mmol) and stirring was continued at room temperature in the dark for 18 h. Purification by reversed phase chromatography (Combiflash, C18 20 g, 5 → 40 % $CH_3CN$ in 0.1 % aq. $NH_3$) gave the intermediate *tert*-butyl ester product as a brown solid after concentration. The intermediate was immediately stirred in TFA (1.5 mL) for 1 h. The mixture was concentrated and residual TFA removed by azeotrope with dichloromethane (2 × 5 mL). 7 N $NH_3$ in MeOH (0.5 mL) and 0.1 % aqueous ammonia (1.5 mL) were added and purification by reversed-phase chromatography (Combiflash, C18 20 g, 5 → 40 % $CH_3CN$ in 0.1 % aq. $NH_3$) gave the title compound as a pale brown solid (16 mg, 39 %) after freeze-drying.

$^{1}H$ NMR (500 MHz, DMSO-$d_6$) δ 12.59 (s, 1H), 8.63 (s, 1H), 8.26 (dd, *J* = 7.9, 1.5 Hz, 1H), 8.02 (t, *J* = 5.5 Hz, 1H), 7.99 (m, 1H), 7.96 (d, *J* = 8.0 Hz, 1H), 7.89 (m, 1H), 7.83 (t, *J* = 7.5 Hz, 1H), 7.61 (d, *J* = 8.3 Hz, 2H), 7.43 (ddd, *J* = 8.1, 5.1, 2.3 Hz, 1H), 7.38 (m, 1H), 7.23 (t, *J* = 9.0 Hz, 1H), 7.01 (br s, 2H), 6.91 (t, *J* = 6.1 Hz, 1H), 6.64 (d, *J* = 8.5 Hz, 2H), 4.47 (d, *J* = 6.0 Hz, 2H), 4.33 (s, 2H), 4.19 – 4.16 (m, 1H), 3.95 (d, *J* = 5.4 Hz, 1H), 3.90 (d, *J* = 5.3 Hz, 1H), 3.64 (s, 1H), 3.58 (s, 1H), 3.50 (br s, 2H), 3.30 (s, overlap with $H_2O$), 3.20 (m, 1H), 3.16 (s, 1H), 2.23 (t, *J* = 7.9 Hz, 2H), 2.03 – 2.00 (m, 1H), 1.92 – 1.87 (m, 1H).

LCMS (Method B): RT = 0.94 min, >99 % @ 254 nm, ES+ 847.3 $[M+H]^+$.

HRMS: found 847.3061; calculated for $C_{41}H_{40}FN_{12}O_8$ $(M+H)^+$ 847.3076.

**Synthesis of 10c**

**4-{2-Fluoro-5-[(4-oxo-3*H*-phthalazin-1-yl)methyl]benzoyl}-1-[3-(*tert*-butoxycarbonylamino)propionyl]piperazine**

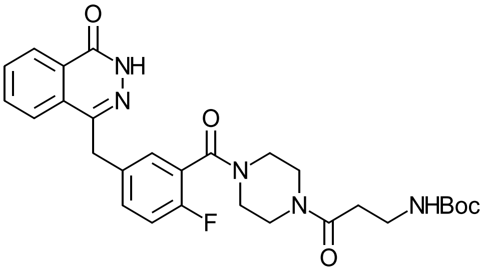


To a stirred solution of compound **7** (200 mg, 0.55 mmol), Boc-β-alanine (125 mg, 0.66 mmol), hydroxybenzotriazole hydrate (38 mg, 0.28 mmol) and triethylamine (167 mg, 230 µL, 1.65 mmol) in DMF (5 mL) was added EDC.HCl (159 mg, 0.83 mmol) and stirring was continued at room temperature for 20 h. The reaction mixture was diluted with ethyl acetate (30 mL) and washed with water (50 mL). The aqueous layer was back-extracted with ethyl acetate (30 mL). Combined organic layers were washed with 2 M hydrochloric acid (30 mL), saturated aqueous $NaHCO_3$ (30 mL), water (2 × 30 mL), brine (20 mL), then dried through a phase separator and concentrated to afford a pale brown gum (120 mg) that was purified by reversed-phase chromatography (Combiflash, C18 20 g, 5 → 40 → 95 % $CH_3CN$ in 0.1 % aq. $NH_3$). Product fractions were combined, and acetonitrile was removed. The resulting aqueous layer was extracted with dichloromethane (2 × 20 mL). Combined organic layers were dried through a phase separator and concentrated to afford the title compound as a colourless solid (75 mg, 25 %) containing amine impurity that was used without further purification.

$^1$H NMR (400 MHz, Chloroform-*d*) δ 9.96 (s, 1H), 8.49 – 8.42 (m, 1H), 7.76 (dd, *J* = 8.2, 4.5 Hz, 2H), 7.71 (d, *J* = 4.2 Hz, 1H), 7.32 (d, *J* = 8.4 Hz, 2H), 7.10 – 7.00 (m, 1H), 4.28 (s, 2H), 3.85 – 3.64 (m, 2H), 3.59 – 3.51 (m, 2H), 3.47 – 3.33 (m, 6H), 3.33 – 3.23 (m, 2H), 1.58 (s, 9H).

LCMS (Method D): RT = 2.35 min, ES+ 560.3 $[M+Na]^+$.

**4-({3-[(4-β-Alanyl-1-piperazinyl)carbonyl]-4-fluorophenyl}methyl)-2*H*-phthalazin-1-one Trifluoroacetate (8c)**

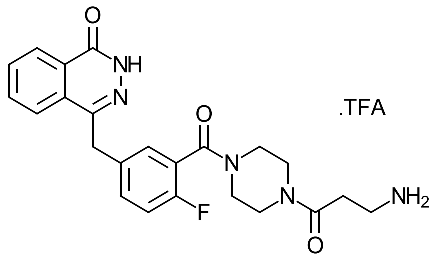


A solution of 4-{2-fluoro-5-[(4-oxo-3*H*-phthalazin-1-yl)methyl]benzoyl}-1-[3-(*tert*-butoxycarbonylamino)propionyl]piperazine (75 mg, 0.14 mmol) was stirred in TFA (2 mL) at room temperature for 1 h, then concentrated and residual TFA removed by azeotrope with dichloromethane (2 × 3 mL). The resulting gum was triturated with diethyl ether (2 × 3 mL, decanted off) and dried to a colourless gum (80 mg, quant.) that was used directly without further purification.

LCMS (Method D): RT = 2.27 min, ES+ 438.3 $(M+H)^+$.

***N*²-(4-(((2-amino-4-oxo-3,4-dihydropteridin-6-yl)methyl)amino)benzoyl)-*N*⁵-(3-(4-(2-fluoro-5-((4-oxo-3,4-dihydrophthalazin-1-yl)methyl)benzoyl)piperazin-1-yl)-3-oxopropyl)-*L*-glutamine (10c)**

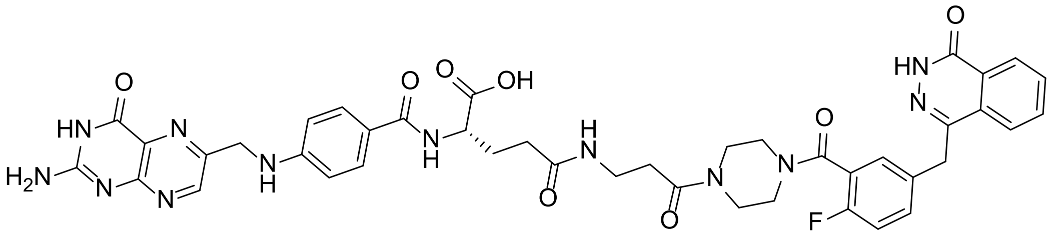

To a stirred mixture of amine trifluoroacetate **8c** (80 mg, 0.14 mmol) and folate active ester **6** (90 mg, 0.15 mmol) in DMSO (1.5 mL) was added triethylamine (99 mg, 136 µL, 0.98 mmol) and stirring was continued at room temperature in the dark for 64 h. Purification by reversed-phase chromatography (Combiflash, C18 20 g, 5 → 50 % $CH_3CN$ in 0.1 % aq. $NH_3$) gave the intermediate *tert*-butyl ester product as a brown solid that was used directly. The intermediate was immediately stirred in TFA (3 mL) for 1 h. The mixture was concentrated and residual TFA was removed by azeotrope with dichloromethane (3 × 4 mL). The resulting gum was triturated with diethyl ether (3 × 4 mL, decanted off) and dried. 7 N $NH_3$ in MeOH (0.5 mL) and 0.1 % aqueous ammonia (1.5 mL) were added and purification by reversed-phase chromatography (Combiflash, C18 20 g, 2 → 30 % $CH_3CN$ in 0.1 % aq. $NH_3$) gave the title compound as a pale brown solid (29 mg, 24 %) after freeze-drying.

$^1$H NMR (500 MHz, DMSO-$d_6$) δ 12.59 (s, 1H), 8.63 (s, 1H), 8.25 (d, *J* = 7.8 Hz, 1H), 7.95 (d, *J* = 8.2 Hz, 2H), 7.88 (t, 2H), 7.82 (t, *J* = 7.5 Hz, 1H), 7.60 (t, 2H), 7.43 (t, *J* = 6.7 Hz, 1H), 7.36 (d, *J* = 6.6 Hz, 1H), 7.23 (t, *J* = 9.0 Hz, 1H), 7.10 (br s, 2H), 6.91 (br s, 1H), 6.63 (t, 2H), 4.49 – 4.44 (m, 2H), 4.32 (s, 2H), 4.18 – 4.14 (m, 1H), 3.62 (s, 1H), 3.56 (s, 1H), 3.52 – 3.45 (m, 2H), 3.36 (s, overlap with $H_2O$), 3.25 –

3.20 (m, 2H), 3.19 – 3.16 (m, 1H), 3.16 – 3.11 (m, 1H), 2.16 – 2.09 (m, 2H), 2.06 – 1.97 (m, 1H), 1.92 – 1.83 (m, 1H).

LCMS (Method B): RT = 0.81 min, 100 % @ 254 nm, ES+ 861.3 $[M+H]^+$.

HRMS: found 861.3234; calculated for $C_{42}H_{42}FN_{12}O_8$ $(M+H)^+$ 861.3226.

**Synthesis of 10d**

**5-(4-{2-Fluoro-5-[(4-oxo-3*H*-phthalazin-1-yl)methyl]benzoyl}-1-piperazinyl)-5-oxovaleric acid**

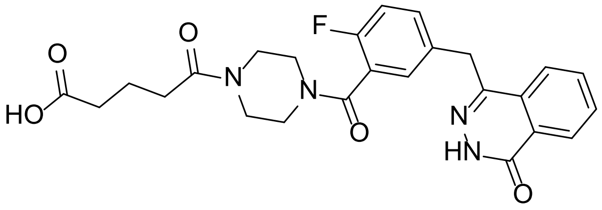

To a stirred solution of compound **7** (185 mg, 0.51 mmol) in dichloromethane (10 mL) was added glutaric anhydride (58 mg, 0.51 mmol) and DIPEA (198 mg, 266 µL, 1.53 mmol), and stirring was continued at room temperature for 2 h. The reaction mixture was diluted with dichloromethane (20 mL) and washed with 0.5 M hydrochloric acid. Layers were separated and the aqueous layer was back-extracted with dichloromethane (30 mL). Combined organic extracts were dried ($MgSO_4$) and concentrated to a colourless gum/solid (200 mg, 82 %).

$^1H$ NMR (500 MHz, Methanol-$d_4$) δ 8.37 (d, *J* = 7.7 Hz, 1H), 7.96 (dd, *J* = 8.1, 3.2 Hz, 1H), 7.92 – 7.81 (m, 2H), 7.49 (tt, *J* = 5.2, 2.7 Hz, 1H), 7.39 (s, 1H), 7.17 (t, *J* = 9.0 Hz, 1H), 4.74 (s, 2H), 4.39 (s, 2H), 3.80 (t, *J* = 5.2 Hz, 1H), 3.74 (t, *J* = 5.1 Hz, 1H), 3.70 – 3.63 (m, 2H), 3.55 – 3.46 (m, 2H), 3.36 – 3.32 (m, 2H), 2.51 (t, *J* = 7.5 Hz, 1H), 2.44 (t, *J* = 7.6 Hz, 1H), 2.37 (q, *J* = 7.6 Hz, 2H), 1.88 (m, 2H).

LCMS (Method C): RT = 2.47 min, ES+ 467.2 $[M+H]^+$.

**tert-Butyl (2-(5-(4-(2-fluoro-5-((4-oxo-3,4-dihydrophthalazin-1-yl)methyl)benzoyl)piperazin-1-yl)-5-oxopentanamido)ethyl)carbamate**

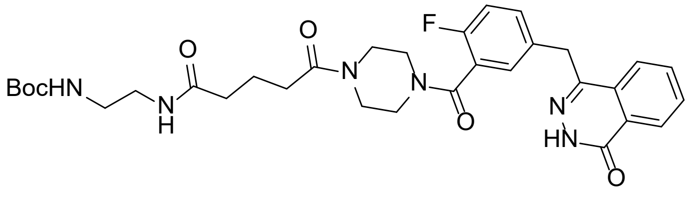

To a stirred solution of 5-(4-{2-fluoro-5-[(4-oxo-3*H*-phthalazin-1-yl)methyl]benzoyl}-1-piperazinyl)-5-oxovaleric acid in dichloromethane (4 mL) was added *N*-Boc-ethylenediamine (99 mg, 98 µL, 0.62 mmol), EDC.HCl (119 mg, 0.62 mmol) and triethylamine (83 mg, 114 µL, 0.82 mmol) and stirring was continued for 24 h. The reaction mixture was diluted with dichloromethane (20 mL), washed with 2 M hydrochloric acid (30 mL), sat. aqueous $NaHCO_3$ (30 mL), water (20 mL), dried through a phase separator and concentrated to afford the title compound as an off-white solid (226 mg, 88 %).

LCMS (Method C): RT = 1.97 min, ES+ 623.4 $[M+H]^+$.

**N-(2-aminoethyl)-5-(4-(2-fluoro-5-((4-oxo-3,4-dihydrophthalazin-1-yl)methyl)benzoyl)piperazin-1-yl)-5-oxopentanamide (8d)**

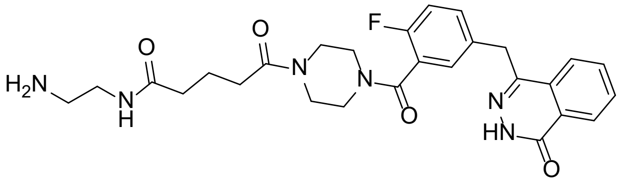

A solution of 4-{2-fluoro-5-[(4-oxo-3*H*-phthalazin-1-yl)methyl]benzoyl}-1-{5-oxo-5-[2-(*tert*-butoxycarbonylamino)ethylamino]valeryl}piperazine (50 mg, 0.08 mmol) was stirred in TFA (1 mL) at room temperature for 1 h. The mixture was concentrated and residual TFA removed by azeotrope with dichloromethane (4 × 2 mL). The resulting gum was triturated with diethyl ether (2 × 2 mL, decanted off) and dried to a colourless gum (52 mg, quantitative).

LCMS (Method C): RT = 1.54 min, ES+ 523.3 $[M+H]^+$.

***N*²-(4-(((2-amino-4-oxo-3,4-dihydropteridin-6-yl)methyl)amino)benzoyl)-*N*⁵-(2-(5-(4-(2-fluoro-5-((4-oxo-3,4-dihydrophthalazin-1-yl)methyl)benzoyl)piperazin-1-yl)-5-oxopentanamido)ethyl)-L-glutamine (10d)**

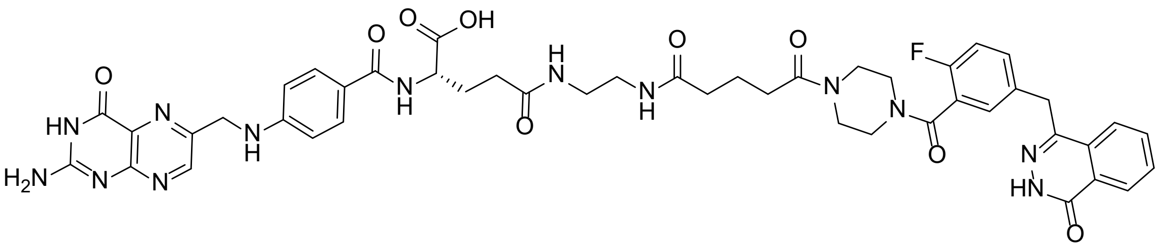

To a stirred mixture of amine trifluoroacetate **8d** (52 mg, 0.08 mmol) and folate active ester **6** (52 mg, 0.09 mmol) in DMSO (1 mL) triethylamine (57 mg, 79 µL, 0.56 mmol) was added and stirring was continued at room temperature in the dark for 18 h. Purification by reversed-phase chromatography

(Combiflash, C18 20 g, 5 → 50 % $CH_3CN$ in 0.1 % aq. $NH_3$) gave the intermediate *tert*-butyl ester product as a brown solid (50 mg, 62 %) after freeze-drying. The intermediate was immediately stirred in TFA (2 mL) for 50 min. The mixture was concentrated and residual TFA removed by azeotrope with dichloromethane (3 × 5 mL). The resulting gum was triturated with diethyl ether (2 × 2 mL, decanted off) and dried. 7 N $NH_3$ in MeOH (0.5 mL) and 0.1 % aqueous ammonia (1.5 mL) were added and purification by reversed-phase chromatography (Combiflash, C18 20 g, 5 → 50 % $CH_3CN$ in 0.1 % aq. $NH_3$) gave the title compound as a pale brown solid (21 mg, 44 %) after freeze-drying.

$^1$H NMR (500 MHz, DMSO-$d_6$) δ 12.60 (s, 1H), 8.63 (s, 1H), 8.26 (dd, *J* = 7.9, 1.4 Hz, 1H), 7.96 (dd, *J* = 8.4, 3.2 Hz, 1H), 7.92 – 7.85 (m, 3H), 7.83 (t, *J* = 7.6 Hz, 1H), 7.60 (dd, *J* = 8.8 Hz, 2H), 7.42 (ddd, *J* = 8.1, 5.1, 2.3 Hz, 1H), 7.36 (d, *J* = 6.5 Hz, 1H), 7.22 (t, *J* = 9.0 Hz, 1H), 7.13 (br s, 2H), 6.90 (t, *J* = 6.2 Hz, 1H), 6.64 (d, *J* = 8.2 Hz, 2H), 4.47 (d, *J* = 6.1 Hz, 2H), 4.32 (s, 2H), 4.15 – 4.12 (m, 1H), 3.64 – 3.60 (m, 1H), 3.58 – 3.54 (m, 1H), 3.53 – 3.47 (m, 2H), 3.38 (s, overlap with $H_2O$), 3.20 – 3.16 (m, 1H), 3.16 – 3.11 (m, 1H), 3.07 – 3.02 (m, 4H), 2.31 (t, *J* = 7.5 Hz, 1H), 2.25 (t, *J* = 7.5 Hz, 1H), 2.16 – 2.05 (m, 4H), 2.02 (q, *J* = 7.2, 6.6 Hz, 1H), 1.91 – 1.83 (m, 1H), 1.73 – 1.65 (m, 2H).

LCMS (Method B): RT = 0.98 min, 100 % @ 254 nm, ES+ 946.3 $[M+H]^+$.

HRMS: found 946.3741; calculated for $C_{46}H_{49}FN_{13}O_9$ $(M+H)^+$ 946.3760.

**Synthesis of 10e**

**2-(3-{2-[4-({2-Fluoro-5-[(4-oxo-3*H*-phthalazin-1-yl)methyl]phenyl}methanoyl)-1-piperazinyl]-2-oxoethylamino}-3-oxopropyloxy)ethylamino 2,2-dimethylpropanoate**

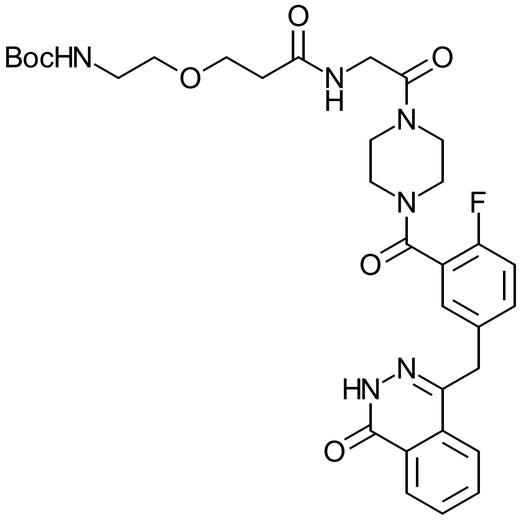

To a stirred solution of **8b** (90 mg, 0.18 mmol, 1 equiv.), 3-(2-((*tert*-butoxycarbonyl)amino)ethoxy)propanoic acid (47 mg, 0.20 mmol, 1.1 equiv.) and Hunig's base (75 mg, 101 µL, 0.58 mmol, 3.2 equiv.) in DMF (2 mL) was added HATU (76 mg, 0.20 mmol, 1.1 equiv.) and stirring was continued for 16 h at room temperature. The reaction mixture was diluted with ethyl acetate (30 mL) and washed with water (50 mL). The aqueous layer was re-extracted with ethyl acetate (30 mL). Combined organic layers were washed with 0.2 M hydrochloric acid (50 mL), water (50 mL), sat. aqueous sodium bicarbonate solution (50 mL), brine (50 mL), dried ($MgSO_4$) and concentrated to afford the title compound as a colourless solid (88 mg, 77 %).

LCMS (Method F): RT = 1.48 min, ES+ 639.3 $[M+H]^+$.

**3-(2-Aminoethyloxy)-1-{2-[4-({2-fluoro-5-[(4-oxo-3*H*-phthalazin-1-yl)methyl]phenyl}methanoyl)-1-piperazinyl]-2-oxoethylamino}-1-propanone trifluoroacetate (8e)**

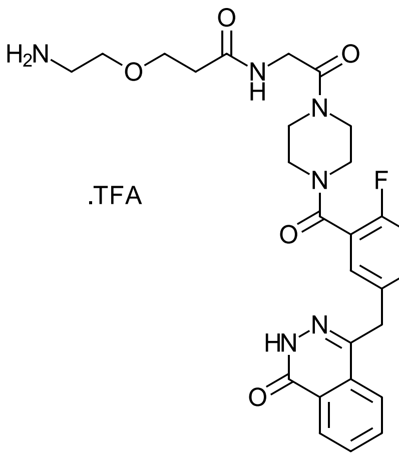


A solution of 2-(3-{2-[4-({2-fluoro-5-[(4-oxo-3*H*-phthalazin-1-yl)methyl]phenyl}methanoyl)-1-piperazinyl]-2-oxoethylamino}-3-oxopropyloxy)ethylamino 2,2-dimethylpropanoate (88 mg, 0.14 mmol) in TFA (1 mL) was stirred at room temperature for 1 h. The reaction mixture was concentrated, co-evaporated with dichloromethane (3 × 3 mL) and triturated with diethyl ether (2 × 3 mL, decanted off) to afford the title compound as a pale brown gum (90 mg, quant.).

$^1$H NMR (500 MHz, DMSO-*d*$_6$) δ 12.59 (s, 1H), 8.26 (d, *J* = 7.8 Hz, 1H), 8.08 (s, 1H), 7.96 (d, *J* = 8.1 Hz, 1H), 7.94 – 7.87 (m, 1H), 7.87 – 7.80 (m, 1H), 7.71 (br s, 3H), 7.49 – 7.42 (m, 1H), 7.38 – 7.34 (m, 1H), 7.25 (t, *J* = 8.4 Hz, 1H), 4.33 (s, 2H), 4.01 – 3.92 (m, 2 H, obscured by water), 3.69 – 3.62 (m, 2H), 3.59 – 3.52 (m, 2H), 3.62 – 3.51 (m, 2H), 3.22 (m, 1H), 3.17 (m, 1H), 3.02 – 2.93 (m, 2H), 2.48 – 2.42 (m, 4H, obscured by DMSO peak).

LCMS (Method F): RT = 1.22 min, 561.2 $[M+Na]^+$.

***N*[2]-(4-(((2-amino-4-oxo-3,4-dihydropteridin-6-yl)methyl)amino)benzoyl)-*N*[5]-(2-(3-((2-(4-(2-fluoro-5-((4-oxo-3,4-dihydrophthalazin-1-yl)methyl)benzoyl)piperazin-1-yl)-2-oxoethyl)amino)-3-oxopropoxy)ethyl)-L-glutamine (10e)**

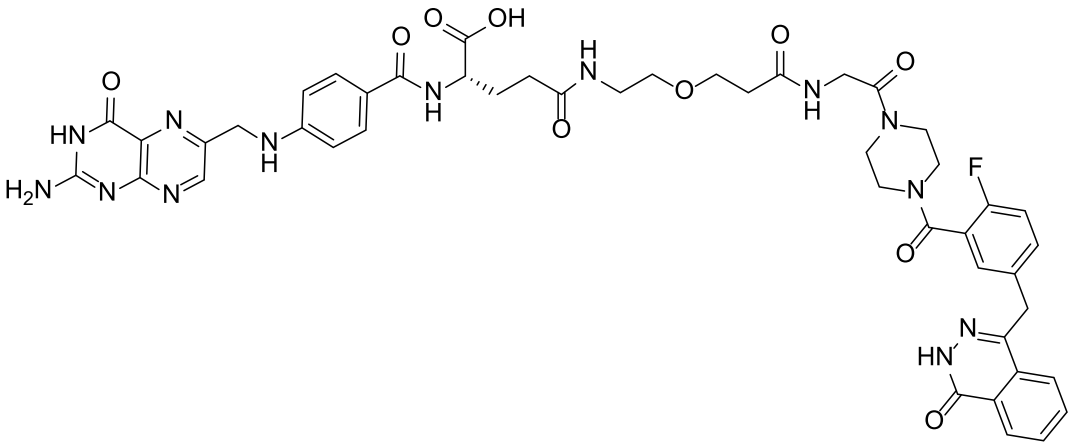

To a stirred mixture of 3-(2-aminoethyloxy)-1-{2-[4-({2-fluoro-5-[(4-oxo-3*H*-phthalazin-1-yl)methyl]phenyl}methanoyl)-1-piperazinyl]-2-oxoethylamino}-1-propanone trifluoroacetate (90 mg, 0.14 mmol, 1 equiv.) and folate active ester **6** (101 mg, 0.17 mmol, 1.2 equiv.) in DMSO (1 mL) was added triethylamine (99 mg, 136 µL, 0.98 mmol, 7 equiv.) and stirring was continued at room temperature in the absence of light for 64 h. Purification by reversed-phase chromatography (Combiflash C18 32 g, 5 → 55 % acetonitrile in aq. 0.1 % ammonia solution). Product fractions were concentrated to afford the *tert*-butyl ester protected product (110 mg). The intermediate was stirred in TFA (2 mL) for 30 min. The reaction mixture was concentrated, co-evaporated with dichloromethane (5 × 2 mL) and triturated with diethyl ether (2 × 2 mL, decanted off) to afford the crude product as a brown solid. Purification by reversed-phase chromatography (Combiflash C18 32 g, 2 → 35 % acetonitrile in aq. 0.1 % ammonia solution) and freeze-drying afforded the title compound as a pale brown solid (59 mg, 44 %).

$^1$H NMR (500 MHz, DMSO-$d_6$) δ 12.59 (s, 1H), 8.62 (s, 1H), 8.25 (d, *J* = 7.8 Hz, 1H), 8.03 – 7.98 (m, 1H), 7.98 – 7.93 (m, 1H), 7.93 – 7.85 (m, 3H), 7.82 (t, *J* = 7.6 Hz, 1H), 7.60 (d, *J* = 8.4 Hz, 2H), 7.46 – 7.40 (m, 1H), 7.40 – 7.35 (m, 1H), 7.23 (t, *J* = 8.9 Hz, 1H), 7.15 (br s, 2H), 6.90 (t, *J* = 6.1 Hz, 1H), 6.64 (d, *J* = 8.6 Hz, 2H), 4.47 (d, *J* = 5.9 Hz, 2H), 4.32 (s, 2H), 4.18 – 4.10 (m, 1H), 4.01 – 3.96 (m, 1H), 3.95 – 3.91 (m, 1H), 3.64 (s, 1H), 3.60 – 3.53 (m, 3H), 3.53 – 3.45 (m, 2H), 3.41 – 3.31 (m, obscured by water), 3.20 (s, 1H), 3.19 – 3.12 (m, 3H), 2.41 – 2.35 (m, 2H), 2.14 (t, *J* = 7.2 Hz, 2H), 2.06 – 1.96 (m, 1H), 1.92 – 1.81 (m, 1H).

LCMS (Method E): RT = 2.48 min, 99 % @ 254 nm, ES+ 962.4 [M+H]$^+$.

HRMS: found 962.3720; calculated for $C_{46}H_{49}FN_{13}O_{10}$ $(M+H)^+$ 962.3709.

**Synthesis of 10f**

**1-{[1-(1,1-Dimethylethyloxycarbonylamino)cyclopropyl]methanoyl}-4-({2-fluoro-5-[(4-oxo-3*H*-phthalazin-1-yl)methyl]phenyl}methanoyl)piperazine**

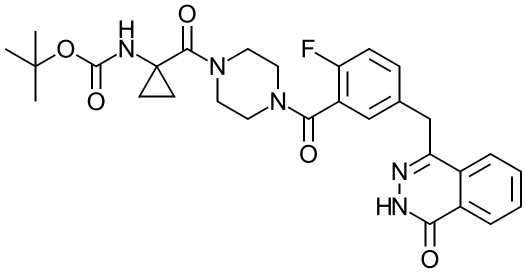

The BOC derivative of **7** (140 mg, 0.30 mmol) was stirred in trifluoroacetic acid (1.0 mL, 1.48 g, 12.98 mmol) for 40 minutes at room temperature before the volatiles were removed. Residual TFA was removed by co-evaporating with DCM, this was repeated until a white foam occurs. This foam was dissolved in DMF (1.5 mL) and 1-(Boc-amino)cyclopropanecarboxylic acid (72 mg, 0.36 mmol), Hünigs base (125 µL, 93 mg, 0.720 mmol) and HATU (137 mg, 0.360 mmol) was added. The resulting solution was left to stir for 2 hours. The reaction was partitioned between ethyl acetate (30 mL) and 0.1 M HCl (20 mL) and the phases separated. The organic phase was washed with water (20 mL), sat. aq. $NaHCO_3$ (20 mL) and then brine (25 mL), dried over $MgSO_4$, filtered and concentrated to yield title compound (182 mg, 98 % yield) as a colourless film which scratches to a white solid.

LCMS (Method H): RT = 1.27 min, ES+ 550.2 $[M+H]^+$.

**1,1-Dimethylethyl (*S*)-2-[(9*H*-fluoren-9-yl)methyloxycarbonylamino]-5-(1-{[4-({2-fluoro-5-[(4-oxo-3*H*-phthalazin-1-yl)methyl]phenyl}methanoyl)-1-piperazinyl]methanoyl}cyclopropylamino)-5-oxopentanoate**

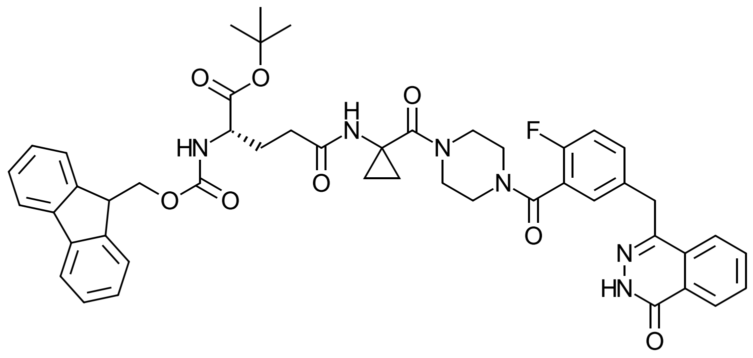

1-{[1-(1,1-Dimethylethyloxycarbonylamino)cyclopropyl]methanoyl}-4-({2-fluoro-5-[(4-oxo-3*H*-phthalazin-1-yl)methyl]phenyl}methanoyl)piperazine (100 mg, 0.182 mmol) was stirred in trifluoroacetic acid (1.0 mL) for 1 hour and the volatiles were removed. Residual trifluoroacetic acid was removed by co-evaporating with dichloromethane several times. The crude residue was dissolved in DMF (1.5 mL) and Fmoc-L-glutamic acid-1-*tert*-butyl ester (93 mg, 0.218 mmol), Hünigs base (76 µL, 56 mg, 0.437 mmol,) and HATU (83.02 mg, 0.218 mmol) was added. The reaction was left to stir at room temperature overnight. The reaction was diluted with EtOAc (40 mL) and washed with 0.2 M HCl (20 mL), brine (20 ml), sat. aqueous $NaHCO_3$ (20 mL) and brine (20 mL), dried over $MgSO_4$, filtered and concentrated to yield title compound (155 mg, 99 % yield) as a white solid.

LCMS (Method H): RT = 1.64 min, ES+ 857.4 $[M+H]^+$.

**1,1-Dimethylethyl(*S*)-2-amino-5-(1-{[4-({2-fluoro-5-[(4-oxo-3*H*-phthalazin-1-yl)methyl]phenyl}methanoyl)-1-piperazinyl]methanoyl}cyclopropylamino)-5-oxopentanoate**

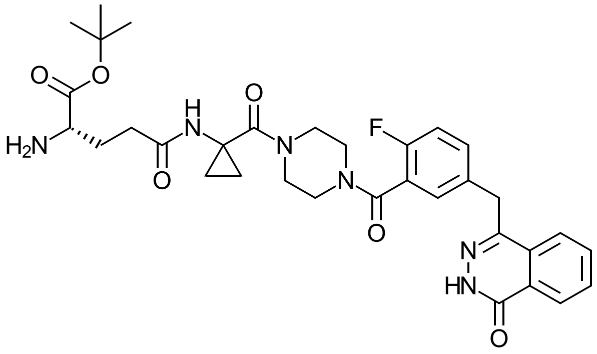

1,1-Dimethylethyl (*S*)-2-[(9*H*-fluoren-9-yl)methyloxycarbonylamino]-5-(1-{[4-({2-fluoro-5-[(4-oxo-3*H*-phthalazin-1-yl)methyl]phenyl}methanoyl)-1-piperazinyl]methanoyl}cyclopropylamino)-5-oxopentanoate (155 mg, 0.180 mmol) was dissolved in a solution of piperidine (715 µL, 616 mg, 7.24 mmol) in acetonitrile (2.9 mL) and the resulting solution was stirred at room temperature for 1 h. The volatiles were removed, and the crude residue was purified by column chromatography (2 → 8% MeOH in DCM) afforded title compound (71 mg, 62 % yield) as a white waxy oil.

$^1$H NMR (500 MHz, Methanol-$d_4$) δ 8.37 (dd, *J* = 7.7, 1.6 Hz, 1H), 7.95 (d, *J* = 7.9 Hz, 1H), 7.88 (td, *J* = 8.1, 7.6, 1.5 Hz, 1H), 7.84 (td, *J* = 7.5, 1.3 Hz, 1H), 7.49 (m, 1H), 7.40 – 7.34 (m, 1H), 7.16 (t, *J* = 9.0 Hz, 1H), 4.39 (s, 2H), 3.74 (br s, 4H), 3.56 (br s, 2H), 2.26 (t, *J* = 7.8 Hz, 2H), 1.95 (m, 1H), 1.79 (m, 1H), 1.46 (s, 9H), 1.39 – 1.24 (m, 2H), 0.98 (d, *J* = 5.8 Hz, 2H).

LCMS (Method H): RT = 1.20 min, ES+ 635.2 $[M+H]^+$.

**$N^2$-(4-(((2-amino-4-oxo-3,4-dihydropteridin-6-yl)methyl)amino)benzoyl)-$N^5$-(1-(4-(2-fluoro-5-((4-oxo-3,4-dihydrophthalazin-1-yl)methyl)benzoyl)piperazine-1-carbonyl)cyclopropyl)-L-glutamine (10f)**

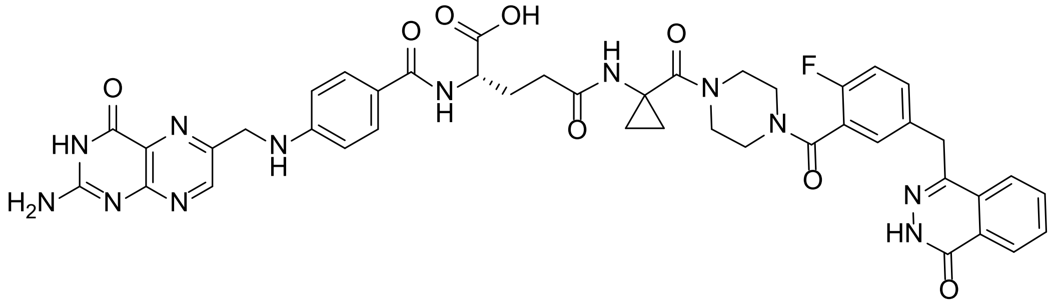


1,1-Dimethylethyl (*S*)-2-amino-5-(1-{[4-({2-fluoro-5-[(4-oxo-3*H*-phthalazin-1-yl)methyl]phenyl}methanoyl)-1-piperazinyl]methanoyl}cyclopropylamino)-5-oxopentanoate (70 mg, 0.110 mmol) was dissolved in DMSO (1.0 mL) and $N^{10}$-(trifluoroacetyl)pteroic acid OSu ester (**4**) (67 mg, 0.132 mmol) and triethylamine (77 µL, 56 mg, 0.551 mmol) were added. The reaction mixture was stirred at room temperature for 2 h in the absence of light. 7N $NH_3$ in methanol (1.5 mL) was added and the resulting solution was stirred for 2 h. The methanolic ammonia was removed and the remaining DMSO solution containing product was purified by reverse phased chromatography (Combiflash C18 32 g, 2 → 35 % acetonitrile in aq. 0.1 % ammonia solution). The fractions were combined and concentrated to provide *tert*-butyl protected product as a yellow solid. The yellow residue was dissolved in TFA (2.0 mL) and stirred at room temperature for 30 minutes. The volatiles were removed, and DCM was added and evaporated several times until a dark yellow solid formed. The product was purified by reverse phased chromatography (Combiflash C18 32 g, 2 → 20 % acetonitrile in aq. 0.1 % ammonia solution). The combined fractions containing product, were purged with $N_2$ and freeze dried to provide the title compound (27 mg, 29% yield over two steps) as a yellow solid.

$^1$H NMR (500 MHz, DMSO-$d_6$) δ 12.62 (s, 1H), 8.64 (s, 1H), 8.52 (br. s, 1H), 8.26 (d, $J$ = 7.8 Hz, 1H), 7.99 (br s, 1H), 7.96 (d, $J$ = 8.0 Hz, 1H), 7.88 (td, $J$ = 7.7, 1.5 Hz, 1H), 7.82 (t, $J$ = 7.8 Hz, 1H), 7.61 (d, $J$ = 8.4 Hz, 2H), 7.46 – 7.39 (m, 1H), 7.38 (d, $J$ = 6.3 Hz, 1H), 7.22 (t, $J$ = 9.0 Hz, 1H), 6.91 (t, $J$ = 6.0 Hz, 1H), 6.63 (d, $J$ = 8.5 Hz, 2H), 4.48 (d, $J$ = 6.0 Hz, 2H), 4.32 (s, 2H), 4.18 (m, 1H), 3.55 (br s, overlap with $H_2O$ peak), 3.14 – 3.07 (m, obscured by water peak), 2.13 (m, 2H), 2.01 (br s, 1H), 1.89 – 1.76 (m, 1H), 1.12 (m, 2H), 0.80 (m, 2H).

LCMS (Method I): RT = 1.75 min, 98 % @ 254 nm, ES+ 873.4 $[M+H]^+$.

HRMS: found 873.3199; calculated for $C_{43}H_{42}FN_{12}O_8$ $(M+H)^+$ 873.3233.

**Synthesis of 10g**

**1-{2-[(*R*)-2-Amino-6-(1,1-dimethylethyloxycarbonylamino)hexanoylamino]ethanoyl}-4-({2-fluoro-5-[(4-oxo-3*H*-phthalazin-1-yl)methyl]phenyl}methanoyl)piperazine (8g)**

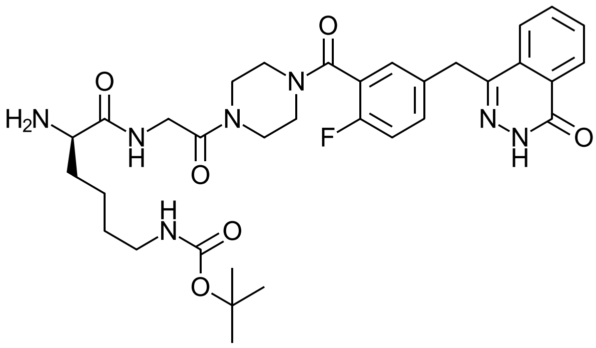


Piperidine (493 µL, 425 mg, 4.99 mmol) was added to a suspension of 9*H*-fluoren-9-yl)methyl *tert*-butyl (6-((2-(4-(2-fluoro-5-((4-oxo-3,4-dihydrophthalazin-1-yl)methyl)benzoyl)piperazin-1-yl)-2-oxoethyl)amino)-6-oxohexane-1,5-diyl)(R)-dicarbamate (prepared via a HATU coupling of Fmoc-D-Lys(Boc)-OH with **8b**) (109 mg, 0.125 mmol) in acetonitrile (1.8 mL). The solids dissolved and the solution was stirred at room temperature for 1 hour. The volatiles were removed under reduced pressure and the pale yellow solid which was dissolved in dichloromethane (20 mL) and washed with water (20 mL), the aqueous phase was extracted with dichloromethane (2 × 10 mL) and the combined organic extracts were dried over MgSO4, filtered and concentrated. The solid was triturated with ether. The title compound (49 mg, 60% yield) was obtained as an off-white solid.

$^1$H NMR (500 MHz, Methanol-$d_4$) δ 8.37 (d, *J* = 7.8 Hz, 1H), 8.00 – 7.91 (m, 1H), 7.92 – 7.79 (m, 2H), 7.53 – 7.45 (m, 1H), 7.41 – 7.36 (m, 1H), 7.16 (t, *J* = 9.0 Hz, 1H), 4.39 (s, 2H), 4.23 – 3.98 (m, 2H), 3.89 – 3.42 (m, 6H), 3.37 (t, *J* = 6.4 Hz, 1H), 3.03 (t, *J* = 6.7 Hz, 2H), 1.81 – 1.64 (m, 2H), 1.65 – 1.52 (m, 2H), 1.51 – 1.38 (m, 13H).

LCMS (Method H): RT = 1.11 min, ES+ 652.3 $[M+H]^+$.

***N*⁵-((*R*)-6-amino-1-((2-(4-(2-fluoro-5-((4-oxo-3,4-dihydrophthalazin-1-yl)methyl)benzoyl)piperazin-1-yl)-2-oxoethyl)amino)-1-oxohexan-2-yl)-*N*²-(4-(((2-amino-4-oxo-3,4-dihydropteridin-6-yl)methyl)amino)benzoyl)-*L*-glutamine (10g)**

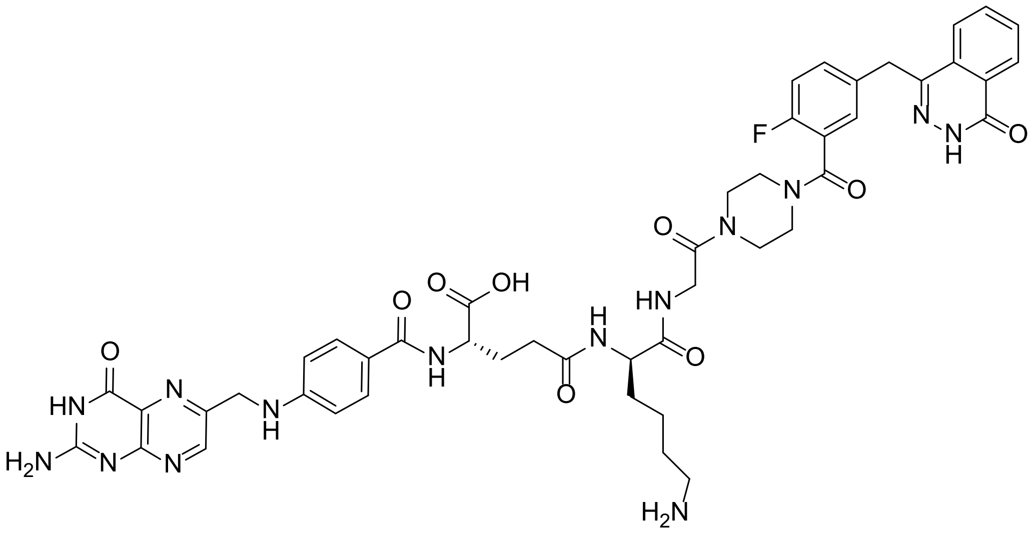


1-{2-[(*R*)-2-Amino-6-(1,1-dimethylethyloxycarbonylamino)hexanoylamino]ethanoyl}-4-({2-fluoro-5-[(4-oxo-3*H*-phthalazin-1-yl)methyl]phenyl}methanoyl)piperazine (50 mg, 0.0767 mmol) and folate active ester **6** (55 mg, 0.092 mmol) were dissolved in DMSO (1.0 mL) and triethylamine (75 µL, 54 mg, 0.537 mmol) was added. The black solution was protected from light and stirred at room temperature overnight. The crude reaction was purified by reversed phased chromatography (Combiflash C18 32 g, 2 → 35 % acetonitrile in aq. 0.1 % ammonia solution), fractions were combined and concentrated by rotary evaporation to yield provide BOC and *tert*-butyl protected product as a yellow solid. The yellow solid was dissolved in TFA (2.0 mL) and stirred for 30 minutes. The reaction was concentrated and residual TFA was co-evaporated with dichloromethane several times to give the crude product as a yellow solid. The crude product was purified by reversed phase chromatography (Combiflash C18 32 g, 2 → 25 % acetonitrile in aq. 0.1 % ammonia solution), and fractions containing product were combined, purged with $N_2$ and freeze dried to yield title compound as a yellow solid (34 mg, 47% over two steps).

$^1$H NMR (500 MHz, DMSO-$d_6$) δ 12.59 (br. s, 1H), 8.63 (s, 1H), 8.25 (d, *J* = 8.1 Hz, 1H), 8.09 (d, *J* = 8.1 Hz, 1H), 7.96 (br s, 1H), 7.94 – 7.85 (m, 2H), 7.83 (t, *J* = 7.5 Hz, 1H), 7.58 (d, *J* = 7.8 Hz, 2H), 7.43 (m, 1H), 7.37 (d, *J* = 6.4 Hz, 1H), 7.23 (t, *J* = 8.9 Hz, 1H), 7.10 (br s, 2H), 6.90 (t, *J* = 6.1 Hz, 1H), 6.64 (d, *J* = 8.5 Hz, 2H), 4.47 (d, *J* = 5.9 Hz, 2H), 4.32 (s, 2H), 4.22 (br s, 1H), 4.13-4.04 (m, 1H), 3.45 – 3.10 (m, overlap with water peak), 2.76 (t, *J* = 6.3 Hz, 2H), 2.23 – 2.10 (m, 2H), 2.04 – 1.94 (m, 1H), 1.93 – 1.83 (m, 1H), 1.72 – 1.61 (m, 1H), 1.56- 1.41 (m, 4H), 1.40 – 1.25 (m, 1H).

LCMS (Method I): RT = 1.73 min, >98 % @ 254 nm, ES+ 975.5 $[M+H]^+$

HRMS: found 975.4011; calculated for $C_{47}H_{52}FN_{14}O_9$ $(M+H)^+$ 975.4026.

**Synthesis of 10h**

**1-[(*R*)-3-*tert*-Butoxy-2-(*tert*-butoxycarbonylamino)propionyl]-4-{2-fluoro-5-[(4-oxo-3*H*-phthalazin-1-yl)methyl]benzoyl}piperazine**

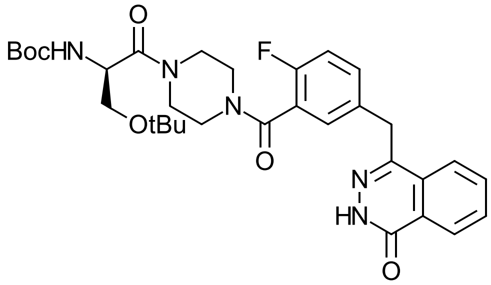


To a stirred solution of 4-({4-fluoro-3-[(1-piperazinyl)carbonyl]phenyl}methyl)-2*H*-phthalazin-1-one (**7**) (150 mg, 0.41 mmol, 1 equiv.), (*R*)-3-*tert*-butoxy-2-((*tert*-butoxycarbonyl)amino)propanoic acid (118 mg, 0.45 mmol, 1.1 equiv.) and Hunig's base (117 mg, 158 µL, 0.90 mmol, 2.2 equiv.) in DMF (3 mL) was added HATU (171 mg, 0.45 mmol, 1.1 equiv.) and stirring was continued for 64 h at room temperature. The reaction mixture was diluted with ethyl acetate (50 mL) and washed with water (50 mL). The aqueous layer was re-extracted with ethyl acetate (50 mL). Combined organic layers were washed with 0.5 M hydrochloric acid (50 mL), brine (50 mL), saturated aqueous sodium bicarbonate solution (50 mL), brine (50 mL), dried ($MgSO_4$) and concentrated to afford the title compound as a pale brown solid (180 mg, 72 %) containing an aliphatic impurity that was used without further purification.

$^1$H NMR (500 MHz, DMSO-$d_6$) δ 12.59 (s, 1H), 8.26 (dd, *J* = 7.8, 1.5 Hz, 1H), 7.97 (d, *J* = 8.0 Hz, 1H), 7.89 (t, *J* = 7.1 Hz, 1H), 7.83 (t, *J* = 7.5 Hz, 1H), 7.47 – 7.41 (m, 1H), 7.40 – 7.35 (m, 1H), 7.24 (t, *J* = 9.0 Hz, 1H), 6.87 – 6.83 (m, 1H), 4.52 – 4.40 (m, 1H), 4.33 (s, 2H), 3.68 – 3.34 (m, 6H), 3.22 (s, 1H), 3.16 (s, 1H), 2.69 (s, 2H), 1.37, 1.34 (2 × s, 9H), 1.10, 1.07 (2 × s, 9H).

LCMS (Method F): RT = 1.83 min, ES+ 610.4 $[M+H]^+$.

**4-{[3-({4-[(*R*)-2-Amino-3-hydroxypropionyl]-1-piperazinyl}carbonyl)-4-fluorophenyl]methyl}-2*H*-phthalazin-1-one (8h)**

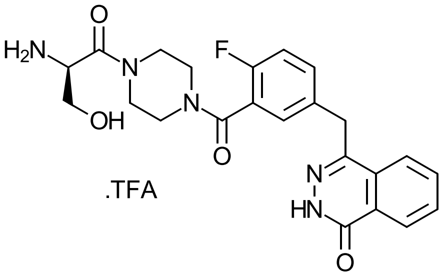


A solution of 1-[(*R*)-3-*tert*-butoxy-2-(*tert*-butoxycarbonylamino)propionyl]-4-{2-fluoro-5-[(4-oxo-3*H*-phthalazin-1-yl)methyl]benzoyl}piperazine (180 mg, 0.30 mmol) in TFA (2 mL) was stirred at room temperature for 1 h. The reaction mixture was concentrated, co-evaporated with dichloromethane (3 × 3 mL). The residue was triturated with diethyl ether (2 × 3 mL, decanted off) and concentrated to afford the title compound as a colourless gum (180 mg, quantitative) that was used without further purification.

LCMS (Method F): RT = 1.14 min, ES+ 454.0 $[M+H]^+$.

***N*$^2$-(4-(((2-amino-4-oxo-3,4-dihydropteridin-6-yl)methyl)amino)benzoyl)-*N*$^5$-((*R*)-1-(4-(2-fluoro-5-((4-oxo-3,4-dihydrophthalazin-1-yl)methyl)benzoyl)piperazin-1-yl)-3-hydroxy-1-oxopropan-2-yl)-*L*-glutamine (10h)**

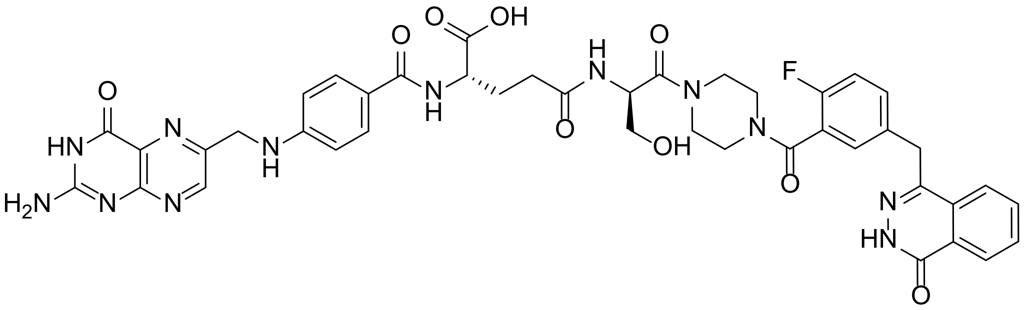

To a stirred mixture of 4-{[3-({4-[(*R*)-2-amino-3-hydroxypropionyl]-1-piperazinyl}carbonyl)-4-fluorophenyl]methyl}-2*H*-phthalazin-1-one (**8h**) (180 mg, 0.30 mmol, 1 equiv.) and the folate active ester **6** (214 mg, 0.36 mmol, 1.2 equiv.) in DMSO (2 mL) was added triethylamine (212 mg, 292 µL, 2.10 mmol, 7 equiv.) and stirring was continued at room temperature in the absence of light for 16 h. Purification by reversed-phase chromatography (Combiflash C18 32 g, 5 → 35 % acetonitrile in aq.0.1 % ammonia solution). Product fractions were concentrated to afford the *tert*-butyl ester protected product (230 mg). The intermediate was stirred in TFA (2 mL) for 1 h. The reaction mixture was concentrated, co-evaporated with dichloromethane (3 × 5 mL) to afford the crude product as a brown solid. Purification by reversed-phase chromatography (Combiflash C18 32 g, 2 → 35 % acetonitrile in aq. 0.1 % ammonia solution) and freeze-drying afforded the title compound as a pale brown solid (94 mg, 36 %).

$^{1}$H NMR (500 MHz, DMSO-$d_6$) δ 12.60 (s, 1H), 8.63 (s, 1H), 8.26 (d, *J* = 7.9 Hz, 1H), 8.07 (d, *J* = 8.1 Hz, 1H), 7.99 – 7.89 (m, 2H), 7.92 – 7.86 (m, 1H), 7.83 (t, *J* = 7.4 Hz, 1H), 7.64 – 7.58 (m, 2H), 7.42 – 7.37 (m, 2H), 7.22 (t, *J* = 8.9 Hz, 1H), 7.09 (br s, 2H), 6.93 (t, *J* = 6.2 Hz, 1H), 6.64 (d, *J* = 8.4 Hz, 2H), 4.91 (br s, 1H), 4.80 and 4.73 (m, 2 × 0.5H), 4.47 (d, *J* = 5.8 Hz, 2H), 4.32 (s, 2H), 4.16 (br s, 1H), 3.68 – 3.36 (m, 8H), 3.20 (s, 1H), 3.15 (s, 1H), 2.21 – 2.17 (m, 2H), 2.02 – 1.98 (m, 1H), 1.89 – 1.85 (m, 1H).

LCMS (Method E): RT = 2.19 min, 100 % at 254 nm, ES+ 877.2 $[M+H]^+$.

HRMS: found 877.3219; calculated for $C_{42}H_{42}FN_{12}O_9$ $(M+H)^+$ 877.3182.

**Synthesis of 10i**

**1-[(*R*)-2-(1,1-Dimethylethyloxycarbonylamino)propanoyl]-4-({2-fluoro-5-[(4-oxo-3*H*-phthalazin-1-yl)methyl]phenyl}methanoyl)piperazine**

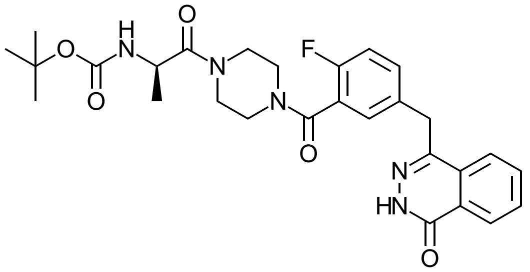

*tert*-Butyl 4-(2-fluoro-5-((4-oxo-3,4-dihydrophthalazin-1-yl)methyl)benzoyl)piperazine-1-carboxylate (47 mg, 0.100 mmol) was dissolved in TFA (1.0 mL) and stirred for 30 minutes at room temperature. TFA was removed by rotary evaporation and co-evaporation with DCM several times. The residue was dissolved in DMF (1.0 mL) and Boc-D-Alanine (23 mg, 0.120 mmol), Hunig's base (42 µL, 31 mg, 0.240 mmol) and HATU (46 mg, 0.120 mmol) was added, and the reaction left to stir at room temperature for 1 hour. EtOAc (20 mL) was added and the reaction washed with 0.2 M HCl (10 mL), $H_2O$, (10 mL) sat. aqueous sodium bicarbonate solution (10 mL) and brine (15 mL). The organic phase was dried over $MgSO_4$, filtered, and concentrated to yield title compound (54 mgs, quant. yield) as a white solid.

LCMS (method H): RT = 1.32 min, ES+ 538.1 $[M+H]^+$.

***N*²-(4-(((2-amino-4-oxo-3,4-dihydropteridin-6-yl)methyl)amino)benzoyl)-*N*⁵-((*R*)-1-(4-(2-fluoro-5-((4-oxo-3,4-dihydrophthalazin-1-yl)methyl)benzoyl)piperazin-1-yl)-1-oxopropan-2-yl)-*L*-glutamine (10i)**

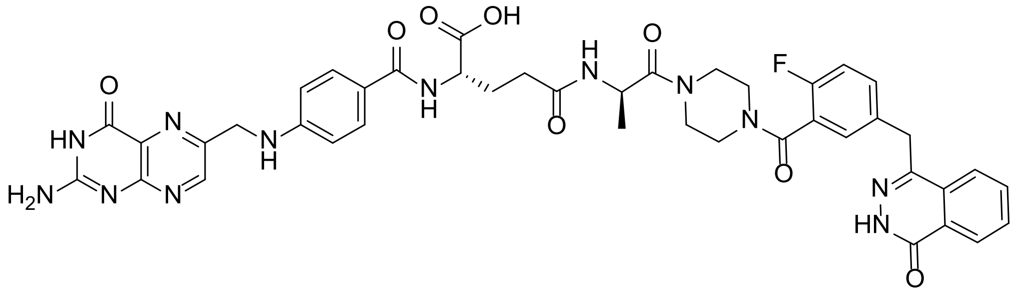

1-[(*R*)-2-(1,1-Dimethylethyloxycarbonylamino)propanoyl]-4-({2-fluoro-5-[(4-oxo-3*H*-phthalazin-1-yl)methyl]phenyl}methanoyl)piperazine 53 mg, 0.100 mmol) was stirred in TFA (1.0 mL) for 1 hour, TFA was removed by rotary evaporation and then by co-evaporating with dichloromethane several times to yield a clear film. LCMS: RT = 1.06 min, ES+ 438.1 $[M+H]^+$. The residue and the folate active ester **6** (71 mg, 0.120 mmol) were dissolved in DMSO (1.0 mL) and triethylamine (98 µL, 71 mg, 0.700 mmol) was added. The dark solution was protected from light and stirred at room temperature overnight. The crude reaction was purified by reversed phased chromatography (Combiflash C18 32 g, 2 → 35 % acetonitrile in aq. 0.1 % ammonia solution), fractions were combined and concentrated by rotary evaporation to yield provide *tert*-butyl protected product as a yellow solid. The yellow solid was dissolved in TFA (2.0 mL) and stirred for 30 minutes. The reaction was concentrated and residual TFA was co-evaporated with dichloromethane several times to give the crude product as a yellow solid. The crude product was purified by reversed phase chromatography and chromatography (Combiflash C18 32 g, 2 → 20 % acetonitrile in aq. 0.1 % ammonia solution), and fractions containing product were combined, purged with $N_2$ and freeze dried to yield title compound as a yellow solid (23 mg, 27% yield over two steps).

$^1$H NMR (500 MHz, DMSO-$d_6$) δ 12.59 (s, 1H), 8.63 (s, 1H), 8.25 (d, *J* = 7.8 Hz, 1H), 8.19 (d, *J* = 7.8 Hz, 1H), 8.00 (br t, 1H), 7.92 (br t, 1H), 7.89 (td, *J* = 7.6, 1.5 Hz, 1H), 7.82 (t, *J* = 7.5 Hz, 1H), 7.62 (t, 2H), 7.46 – 7.36 (m, 2H), 7.22 (t, *J* = 9.0 Hz, 1H), 7.02 (br s, 2H), 6.91 (bt t, *J* = 6.1 Hz, 1H), 6.64 (d, *J* = 8.3 Hz, 2H), 4.71 – 4.59 (m, 1H), 4.47 (br s, 2H), 4.32 (s, 2H), 4.22 (br s, 1H), 3.67 (s, overlap with $H_2O$), 3.57 (d, *J* = 6.2 Hz, overlap with $H_2O$), 3.20 (s, 1H), 3.17 – 3.08 (m, 1H), 2.24 – 2.13 (m, 2H), 2.06 – 1.97 (m, 1H), 1.93 – 1.81 (m, 1H), 1.11, 1.08 (2 x d, *J* = 6.8 Hz, 3H).

LCMS (Method I): RT = 1.68 min, >98 % @ 254 nm, ES+ 861.1 $[M+H]^+$.

HRMS: found 861.3200; calculated for $C_{42}H_{42}FN_{12}O_8$ $(M+H)^+$ 861.3233.

**Synthesis of 11**

**(*R*)-4-(4-(((2-amino-4-oxo-3,4-dihydropteridin-6-yl)methyl)amino)benzamido)-5-(*tert*-butoxy)-5-oxopentanoic acid**

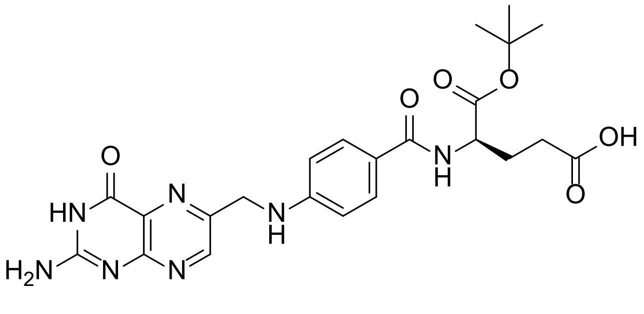

To a stirred mixture of $N^{10}$-trifluoroacetate pteroic acid NHS-activated ester **4** (200 mg, 0.40 mmol) and *D*-glutamic acid α-*tert*-butyl ester (163 mg, 0.80 mmol) in DMF (5 mL) was added triethylamine

(81 mg, 112 µL, 0.80 mmol) and stirring was continued at room temperature in the dark for 18 h. LCMS showed full conversion to the intermediate amide ($(M+H)^+$= 594.2). Ammonia (7 N) in methanol (2 mL) was added and stirring continued for 1 h. The reaction mixture was concentrated to a brown solid that stirred in water (15 mL) for 30 min. The resulting solid was collected by suction filtration, washed with water (3 × 30 mL) and dried under vacuum to afford the title compound as a brown solid (150 mg, 75 %).

$^1$H NMR (500 MHz, DMSO-$d_6$) δ 11.53 (s, 3H), 8.64 (s, 1H), 8.13 (d, *J* = 7.6 Hz, 1H), 7.64 (d, *J* = 8.3 Hz, 2H), 6.93 (t, *J* = 6.1 Hz, 1H), 6.86 (br s, 2 H), 6.64 (d, *J* = 8.4 Hz, 2H), 4.48 (d, *J* = 6.0 Hz, 2H), 4.26 (td, *J* = 8.8, 5.4 Hz, 1H), 2.31 (t, *J* = 7.5 Hz, 2H), 2.00 (dq, *J* = 13.3, 6.9 Hz, 1H), 1.89 (ddd, *J* = 16.7, 14.2, 7.7 Hz, 1H), 1.39 (s, 9H).

LCMS (Method A): RT = 1.30 min, ES+ 498.2 $[M+H]^+$.

**1-(*tert*-butyl)5-(2,5-dioxopyrrolidin-1-yl)(4-(((2-amino-4-oxo-3,4-dihydropteridin-6-yl)methyl)amino)benzoyl)-*D*-glutamate**

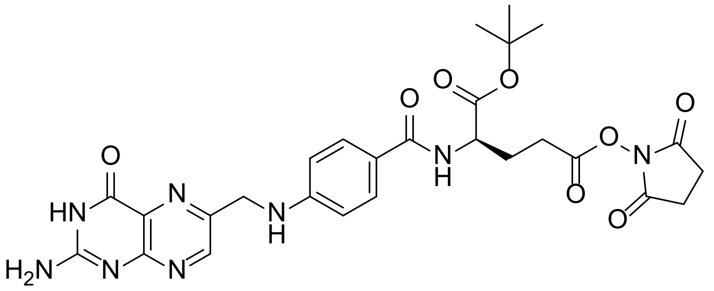


To a stirred, cooled mixture of (*R*)-4-(4-(((2-amino-4-oxo-3,4-dihydropteridin-6-yl)methyl)amino)benzamido)-5-(*tert*-butoxy)-5-oxopentanoic acid (150 mg, 0.30 mmol), *N*-hydroxysuccinimide (69 mg, 0.60 mmol), and 4-dimethylaminopyridine (2 mg, 0.02 mmol) in DMF (4.7 mL) at 0 °C was added *N,N'*-diisopropylcarbodiimide (51 mg, 62 µL, 0.40 mmol) and stirring was continued with warming to room temperature over 40 h. The reaction mixture was concentrated to a dark brown solid that was stirred in dichloromethane-diethyl ether (3:1, 8 mL) for 1 h. The resulting precipitate was collected by suction filtration and washed with diethyl ether (3 × 20 mL). The solid was slurried in dichloromethane (2 × 20 mL), collected by suction filtration, washed with diethyl ether (2 × 15 mL) and dried to afford the title compound as a dark brown solid (150 mg, 84 %) at 90 % purity that was used without further purification.

$^{1}H$ NMR (500 MHz, DMSO-$d_6$) δ 11.43 (s, 1H), 8.65 (s, 1H), 8.21 (d, *J* = 7.5 Hz, 1H), 7.65 (d, *J* = 8.3 Hz, 2H), 6.95 (t, *J* = 6.1 Hz, 1H), 6.88 (s, 2H), 6.65 (d, *J* = 8.3 Hz, 2H), 4.48 (d, *J* = 6.1 Hz, 2H), 4.32 (td, *J* = 8.4, 7.8, 5.0 Hz, 1H), 2.80 (s, 4H), 2.86 – 2.69 (m, 2H), 2.15 – 1.99 (m, 2H), 1.39 (s, 9H).

LCMS (Method A): RT = 1.43 min, ES+ 595.3 $[M+H]^+$.

**$N^2$-(4-(((2-amino-4-oxo-3,4-dihydropteridin-6-yl)methyl)amino)benzoyl)-$N^5$-(2-(4-(2-fluoro-5-((4-oxo-3,4-dihydrophthalazin-1-yl)methyl)benzoyl)piperazin-1-yl)-2-oxoethyl)-*D*-glutamine (11)**

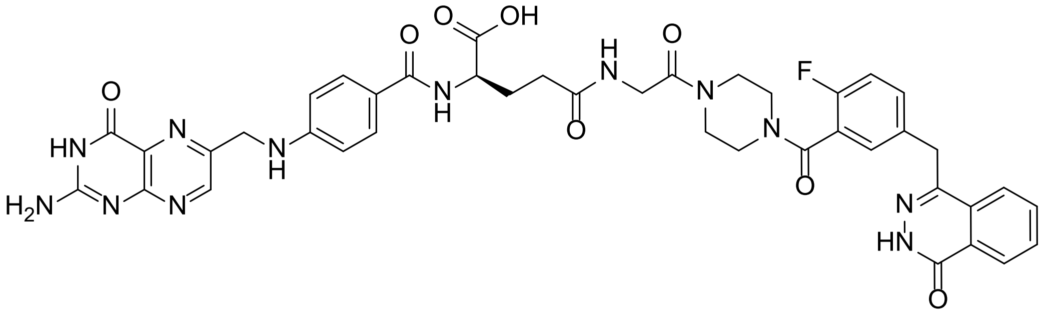

To a stirred mixture of 4-({4-fluoro-3-[(4-glycyl-1-piperazinyl)carbonyl]phenyl}methyl)-2*H*-phthalazin-1-one trifluoroacetate (90 mg, 0.15 mmol) and 1-(*tert*-butyl) 5-(2,5-dioxopyrrolidin-1-yl) (4-(((2-amino-4-oxo-3,4-dihydropteridin-6-yl)methyl)amino)benzoyl)-*D*-glutamate (101 mg, 0.17 mmol) in DMSO (2 mL) was added triethylamine (106 mg, 146 µL, 1.05 mmol) and stirring was continued at room temperature in the dark for 16 h. Purification by reversed-phase chromatography (Combiflash, C18 20 g, 5 → 50 % $CH_3CN$ in 0.1 % aq. $NH_3$) gave the intermediate *tert*-butyl ester product as a brown solid after concentration. The intermediate was immediately stirred in TFA (2 mL) for 1 h. The mixture was concentrated and residual TFA removed by azeotrope with dichloromethane (2 × 5 mL). 7 N $NH_3$ in MeOH (0.5 mL) and 0.1 % aqueous ammonia (1.5 mL) was added and purification by reversed-phase chromatography (Combiflash, C18 20 g, 5 → 35 % $CH_3CN$ in 0.1 % aq. $NH_3$) gave the title compound as a pale brown solid (71 mg, 56 %) after freeze-drying.

$^{1}H$ NMR (500 MHz, DMSO-$d_6$) δ 12.59 (s, 1H), 8.63 (s, 1H), 8.26 (d, *J* = 7.8 Hz, 1H), 8.01 (t, *J* = 5.6 Hz, 1H), 7.96 (d, *J* = 8.1 Hz, 1H), 7.89 (dd, *J* = 12.9, 7.4 Hz, 2H), 7.83 (t, *J* = 7.5 Hz, 1H), 7.61 (d, *J* = 8.3 Hz, 2H), 7.46 – 7.39 (m, 1H), 7.38 (s, 1H), 7.23 (t, *J* = 9.0 Hz, 1H), 7.12 (br s, 2H), 6.91 (t, *J* = 6.2 Hz, 1H), 6.64 (d, *J* = 8.4 Hz, 2H), 4.47 (d, *J* = 6.0 Hz, 2H), 4.32 (s, 2H), 4.20 – 4.14 (m, 1H), 3.97 – 3.93 (m, 1H), 3.92 – 3.87 (m, 1H), 3.65 – 3.62 (m, 1H), 3.60 – 3.57 (m, 1H), 3.51 – 3.48 (m, 2H), 3.40 – 3.33 (m, overlap with $H_2O$), 3.22 – 3.18 (m, 1H), 3.17 – 3.14 (m, 1H), 2.22 (t, *J* = 7.9 Hz, 2H), 2.05 – 2.00 (m, 1H), 1.91 – 1.85 (m, 1H).

LCMS (Method B): RT = 0.93 min, 100 % at 254 nm, ES+ 847.3 $[M+H]^+$.

HRMS: found 847.3061; calculated for $C_{41}H_{40}FN_{12}O_8$ $(M+H)^+$ 847.3076.
